# BEdeepon: an in silico tool for prediction of base editor efficiencies and outcomes

**DOI:** 10.1101/2021.03.14.435303

**Authors:** Chengdong Zhang, Zimeng Yu, Daqi Wang, Tao Qi, Yuening Zhang, Linghui Hou, Feng Lan, Jingcheng Yang, Leming Shi, Sang-Ging Ong, Hongyan Wang, Yongming Wang

**Author notes:** Equal contributors (Chengdong Zhang, Zimeng Yu, Daqi Wang, Tao Qi).

## Abstract

Base editors enable direct conversion of one target base into another in a programmable manner, but conversion efficiencies vary dramatically among different targets. Here, we performed a high-throughput gRNA-target library screening to measure conversion efficiencies and outcome product frequencies at integrated genomic targets and obtained datasets of 60,615 and 73,303 targets for ABE and CBE, respectively. We used the datasets to train deep learning models, resulting in ABEdeepon and CBEdeepon which can predict on-target efficiencies and outcome sequence frequencies. The software is freely accessible via online web server http://www.deephf.com/#/bedeep/bedeepon.

## Introduction

Base editors are fusion of catalytically impaired Cas9 nuclease and adenosine deaminase (ABE) or cytosine deaminase (CBE) that introduce desired point mutations in the target region enabling precise editing of genomes ^1-5^. Base editors have been successfully used in diverse organisms including prokaryotes, plants, fish, frogs, mammals and human embryos ^6-9^. Though simple in concept, the success of base editing depends on the choice of the target sequence. First, efficiency of base conversion varies dramatically among different target sequences, and users need to select a target sequence with high efficiency. Second, there are often more than one editable nucleotide in the editing window and unwanted concurrent mutations should be avoided. Experimental evaluation of a target sequence is time-consuming, prompting us to develop in silico tools for target sequence evaluation.

We have previously established a library containing over 80,000 gRNA-target sequence pairs, covering ∼20,000 human genes ^10^. In this study, we made use of this library to screen both base editors and obtained the outcome product frequencies of 60,615 and 73,303 target sequences for ABE and CBE, respectively. The resulting outcomes were used to train a deep learning model, resulting in ABEdeepon and CBEdeepon which can predict efficiency and outcome frequency distributions of ABE and CBE, respectively.

These models were accessible via http://www.deephf.com/#/bedeep, which will greatly facilitate base editor application.

## Results

### Guide RNA-target pair strategy for test of conversion efficiency and outcomes

A previously generated library ^10^ containing over 80,000 gRNA-target sequence pairs was used for ABE and CBE screening. Optimized version of ABE (ABEmax) and CBE (AncBE4max)^11^ was used in this study. Because of the large size, we used the Sleeping Beauty (SB) transposon ^12-14^ to integrate each base editor into the genome and generated single cell-derived cell lines. Next, we evaluated the nucleotide conversion efficiency with three targets at different time points. As expected, the efficiency increased over time for all three targets (Supplementary Fig. 1). One target displayed ∼100% gene conversion at day

We selected day 5 to measure editing efficiency for the library. We packaged the gRNA-target library into lentiviruses and transduced them into recipient cells. Five days after transduction, genomic DNA was extracted, and synthesized targets were PCR-amplified for deep-sequencing (Fig.1a). The reads containing canonical base editing (A to G for ABE, C to T for CBE) and unedited reads were used for outcome frequency distribution. A target efficiency can be calculated by 1 subtract unedited outcome frequency (see **Data analysis** section of **Methods** part). The screening assay was experimentally repeated twice, and conversion efficiency in two independent replicates showed high correlation for both ABE and CBE (Spearman correlation, 0.891 for ABE and 0.934 for CBE). Data from the two replicates were combined together for subsequent analysis. We obtained valid efficiencies (reads number>100) of 60,615 targets with 5,761,833 outcomes for ABE and 73,303 targets with 5,513,919 outcomes for CBE.

**Figure 1.**
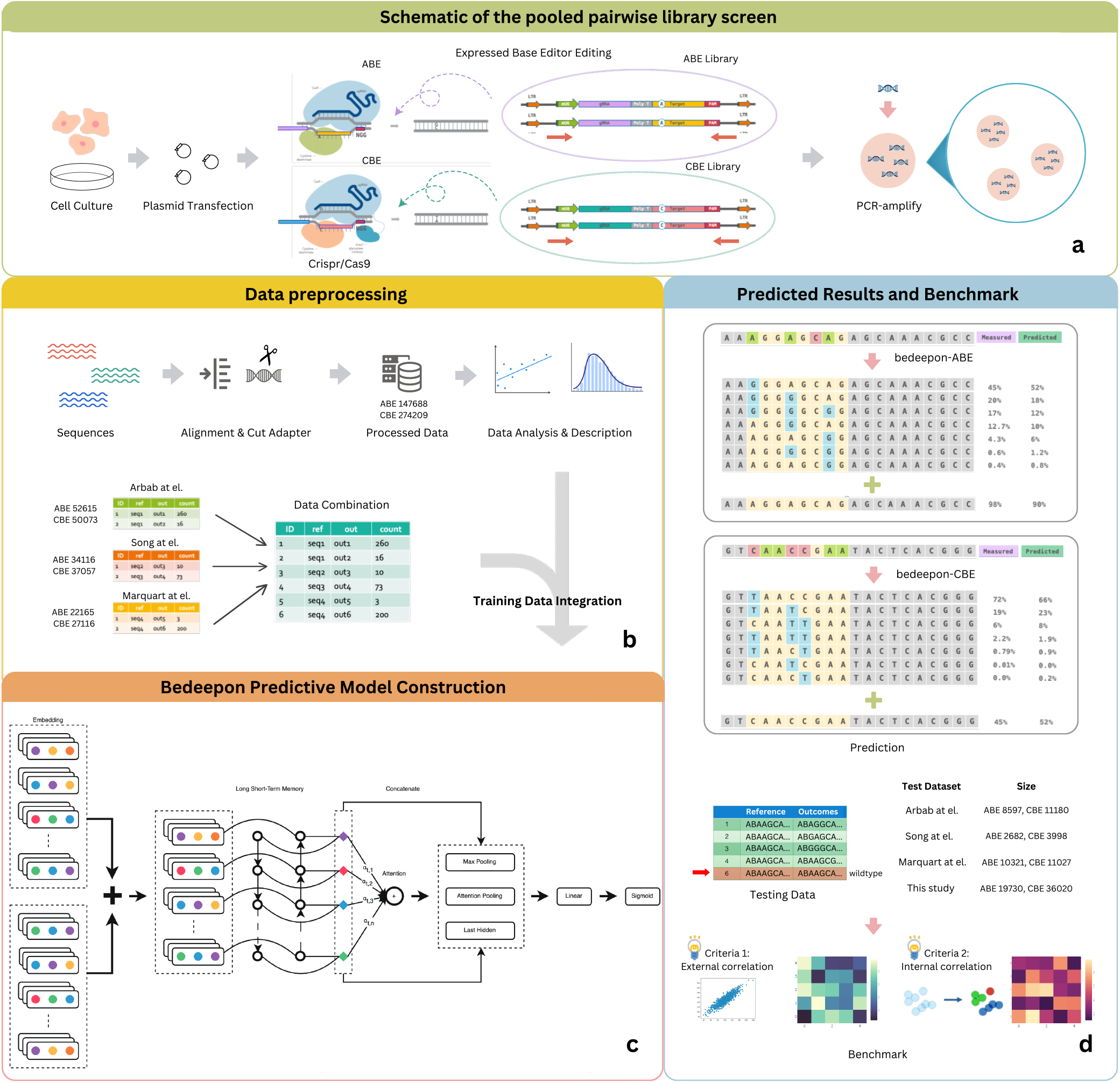
The general workflow of bedeepon. (**a**) Schematic of the pooled pairwise library screen. A library of 80,263 gRNA-target pairs was packed into viruses and transduced into cells expressing ABE or CBE base editors. The integrated target sites were PCR-amplified for deep-sequencing analysis. Red arrows indicate primers for target site amplification; stars indicate nucleotide conversion. (**b**) Procedure of preprocessing data and combination. After we got totally valid (reads number>100) outcomes from our experiment and analysis, we split our training datasets with 164,129 ABE outcomes and 304,866 CBE outcomes, they are combined with 52,615 (Arbab et al.), 34,116 (Song et al.) and 22,165 (Marquart at el.) of ABE outcomes; 50,073 (Arbab et al.), 37,057 (Song et al.) and 27,116 (Marquart at el.) of CBE outcomes to generate an integral training dataset.(**c**) Constructing a predictive bedeepon model with BiLSTM network using sequencing data. (**d**) Generating predicted editing results with model and evaluating performance in four testing datasets. Deep-sequencing results overall outcomes that revealed that A to G conversion (green) for ABE and C to T (red) conversion for CBE occurred, the edited bases are annoted with blue. Then benchmarking on four testing datasets from Arbab et al., Song et al., Marquart at el. and datasets splited from this study (train : test = 9 : 1) on two different criterias, external correlation and internal correlation.

Sequencing results revealed that A to G conversion for ABE and C to T conversion for CBE occurred (Fig. 1b). Although infrequent, C to non-C conversion for CBE and A to non-A conversion for ABE could also be observed (Fig. 1b). The ABE products (purity ranging from 98.43% to 99.69%) were purer than CBE (purity ranging from 95.55% to 97.78%, Supplementary Fig. 2), consistent with a previous report ^2^. We also observed indels with mean efficiency of 0.13 for ABE and mean efficiency of 0.16 for CBE, whereas mean frequency of background indels in the plasmid library was 0.06.

**Figure 2.**
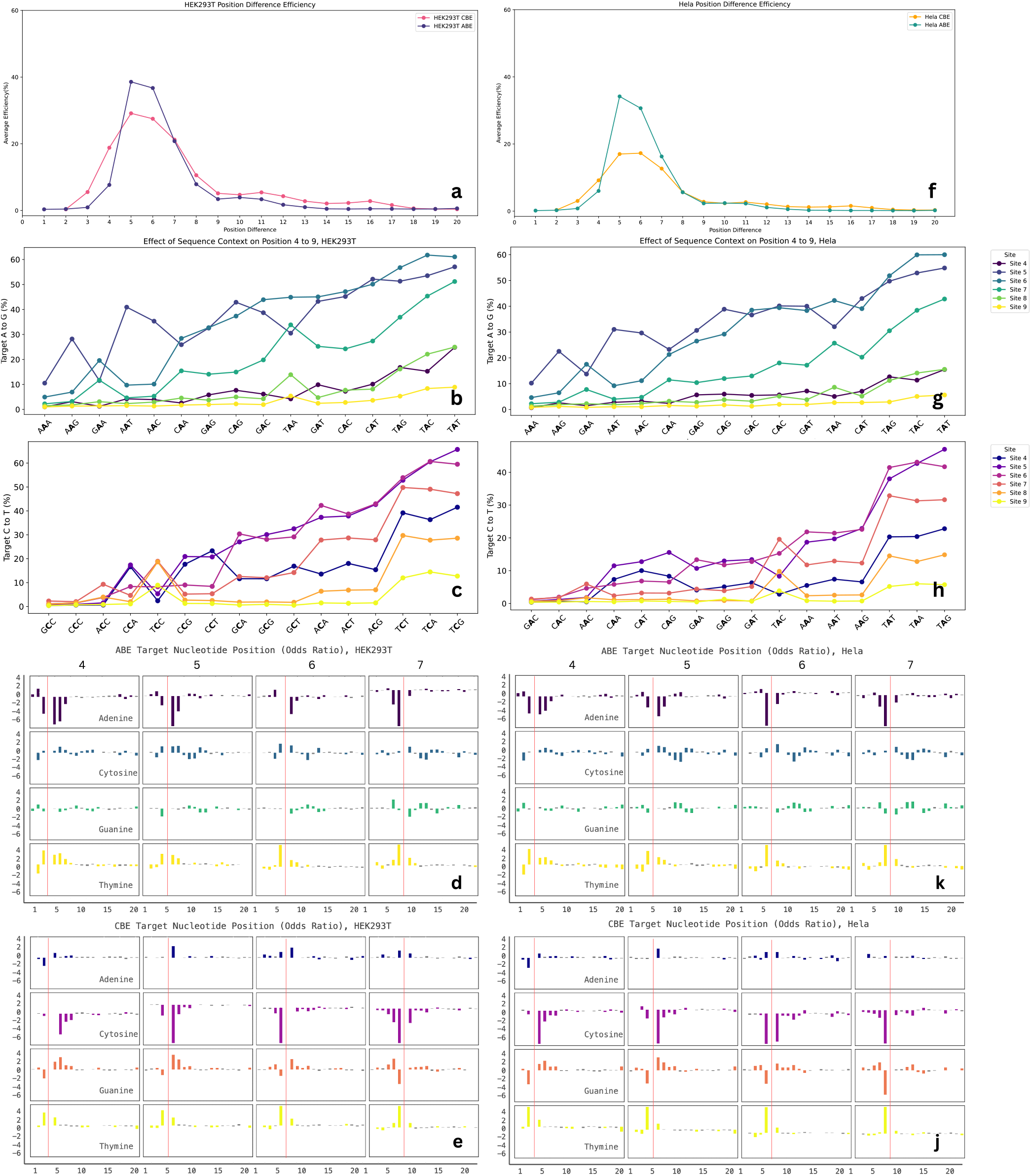
**The relationship between editing efficiency and base position or sequence** context, and conversion efficiency influenced by nucleotides on the target sequence in a high-throughput manner. (**a,f**) C to T or A to G conversion efficiency at different single base position, HEK293T and Hela cell. (**b,c,g,f**) Effect of the sequence context surrounding the target nucleotide (bold A or C) on the conversion efficiency at protospacer positions 4 to 9. Sequence preferences at each position in efficient (top 20%) vs. inefficient (bottom 20%) targets for ABE (**d,k**) and CBE (**e,j**) on position 4 to 7. Nucleotide position on the targets is shown below. The log odds ratios of nucleotide frequencies between efficient and inefficient target sequences are represented on the y axis. Target nucleotide position is indicated by red lines.

### Positional effects on nucleotide conversion efficiency

The large-scale dataset generated here allowed us to analyze the positional effects on nucleotide conversion efficiency. The five most efficient positions for nucleotide conversion are 4-8 for both ABE and CBE with PAM at position 21-23 (Fig. 1c), consistent with previously reported editing windows ^2, 11^. Next, we investigated the influence of nearby nucleotides on conversion efficiency for position 4-9. Both nucleotides surrounding the target A influenced conversion efficiency with the upstream nucleotide having a stronger influence for ABE. The upstream nucleotide preference followed the order T>C>G>A (Fig. 1d). In contrast, only upstream nucleotide surrounding the target C influenced conversion efficiency for CBE following the preference order T>C>A>G (Fig. 1e).

Next, we investigated nucleotide preferences at each position in the target sequences associated with high editing efficiencies. The results revealed that ABE editing efficiencies were strongly influenced by both upstream and downstream nucleotides immediately adjacent to the target nucleotide (target nucleotide, ±1 bp, Fig. 2). The nearby nucleotides A and G have negative influence on editing efficiency, while the nearby T have positive influence on editing efficiency. The upstream C has positive influence on editing efficiency for target nucleotide positions 5 and 6, but has negative influence on target nucleotide positions 4 and 7. The downstream C has positive influence on editing efficiency. CBE editing efficiencies were only strongly influenced by the upstream nucleotide immediately adjacent to the target nucleotide (target nucleotide, +1 bp, Fig. 2). The upstream nucleotides A and G have negative influence on editing efficiency, while the upstream T have positive influence on editing efficiency. Upstream C has positive influence on editing efficiency for target nucleotide positions 5-7, but has negative influence on target nucleotide position 4.

### Correlation of editing efficiency between SpCas9 and base editors

The gRNA-target library in this study was recently used to screen for SpCas9 nuclease ^10^, allowing us to investigate the efficiency correlation between SpCas9 and base editors. We calculated the correlations between SpCas9 nuclease and base editors for each quartile of base editor efficiencies. The correlation (R<0.15) was low for both ABE and CBE (Supplementary Fig. 3a-b). We also calculated the correlations between SpCas9 nuclease and base editors for each quartile of SpCas9 efficiencies. Although the corrections were low, quartile 1 achieved much higher correlation than other quartiles for both ABE and CBE (R=0.21 for ABE, R=0.23 for CBE, Supplementary Fig. 3c-d). We further investigated the relationship between SpCas9 mean efficiencies and base editor mean efficiencies for each quartile of base editor efficiencies. The results revealed that SpCas9 efficiency was similar in each quartile of base editor efficiency (Supplementary Fig. 4a-b). We also investigated the relationship between SpCas9 mean efficiencies and base editor mean efficiencies for each quartile of SpCas9 efficiencies. The results revealed that quartile 2-4 achieved similar mean efficiency, while quartile 1 achieved a little lower mean efficiency (Supplementary Fig. 4c-d). Collectively, these data suggested SpCas9 efficiency and base editor efficiency had a very low correlation for a target.

### Developing models for prediction of on-target efficiencies and outcomes

Next, we developed models for prediction of on-target efficiencies and outcome sequence frequencies. Since a target generally contains multiple editable nucleotides, there might exit multiple editing outcome sequences. We first programmed this conversion scheme to obtain all possible editing outcome sequences, and then assigned the measured editing frequency to each outcome sequence. We used the ABE editing datasets to train a shared embedding based deep learning model. The targets were randomly split to 56,432 (5,390,492 outcomes) for training, 1,152 (98,958 outcomes) for internal validation, and 3,031 (272,383 outcomes) held out for testing. The resulting model was named “ABEdeepon” which can predict editing efficiencies and outcome sequence frequencies (Fig. 3a). ABEdeepon achieved a Spearman correlation of 0.892 (r=0.878, MSE=0.024) for prediction of editing efficiency with testing dataset (Fig. 3b). To evaluate the performance for prediction of outcome frequency distribution, we tested 2,342 targets with at least 4 outcomes and achieved mean Spearman correlation of 0.872 (r=0.920, MSE=0.011, KL divergence=0.003, Fig. 3c). To test whether ABEdeepon works in other cell types, we used the same library to generate datasets in HeLa cells and obtained 58,445 valid targets with 6,292,774 outcomes (reads number≥100). We extracted 2,636 targets present in HeLa testing dataset as external testing dataset. The editing efficiency prediction achieved Spearman correlation of 0.886 (r=0.830, MSE=0.024) and the outcome frequency distribution prediction achieved mean Spearman correlation of 0.857 (r=0.883, MSE=0.020, KL divergence=0.004, Supplementary Fig. 5a-b).

**Figure 3.**
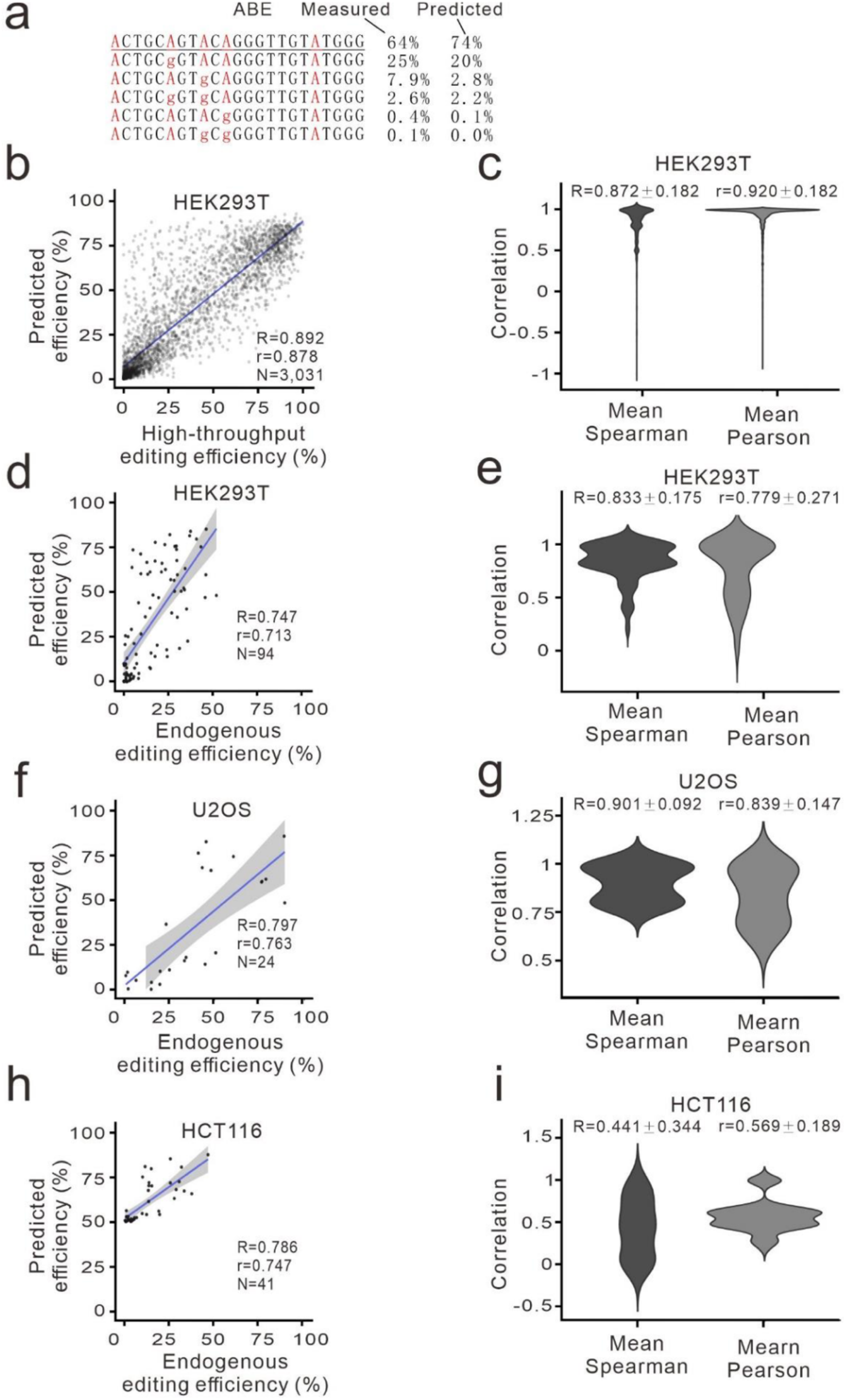
Evaluation of ABEdeepon for editing efficiency and outcome prediction. (**a**) Example of ABEdeepon prediction for a given target. Measured efficiencies are listed for comparison. Original target sequence is underlined. (**b**) Evaluation of ABEdeepon for conversion efficiency with testing dataset. (**c**) Evaluation of ABEdeepon for conversion outcome sequence frequencies with testing dataset. (**d, f, h)** Evaluation of ABEdeepon for conversion efficiency with endogenous editing dataset generated in HEK293T, U2OS and HCT116 cells, respectively. (**e, g, i**) Evaluation of ABEdeepon for conversion outcome sequence frequencies with endogenous editing dataset generated in HEK293T, U2OS and HCT116 cells, respectively.

To evaluate the performance of ABEdeepon for endogenous targets, we collected endogenous editing data generated in HEK293T, U2OS and HCT116 cells from literatures ^15^ and evaluated these datasets with our model, achieving high correlation for efficiencies (Spearman correlation: 0.747-0.797) and mild or high correlation for outcomes (mean Spearman correlation: 0.441-0.901, Fig 4 d-i). To evaluate the performance of the ABEdeepon in induced pluripotent stem cells (iPSCs), we generated dataset for 23 endogenous targets in human iPSCs. ABEdeepon achieved weak Spearman correlation of 0.227 for efficiency prediction probably due to the very low base editing efficiency in iPSCs, and mild mean Spearman correlation of 0.574 for outcome frequency prediction (Supplementary Fig. 5c-d).

**Figure 4.**
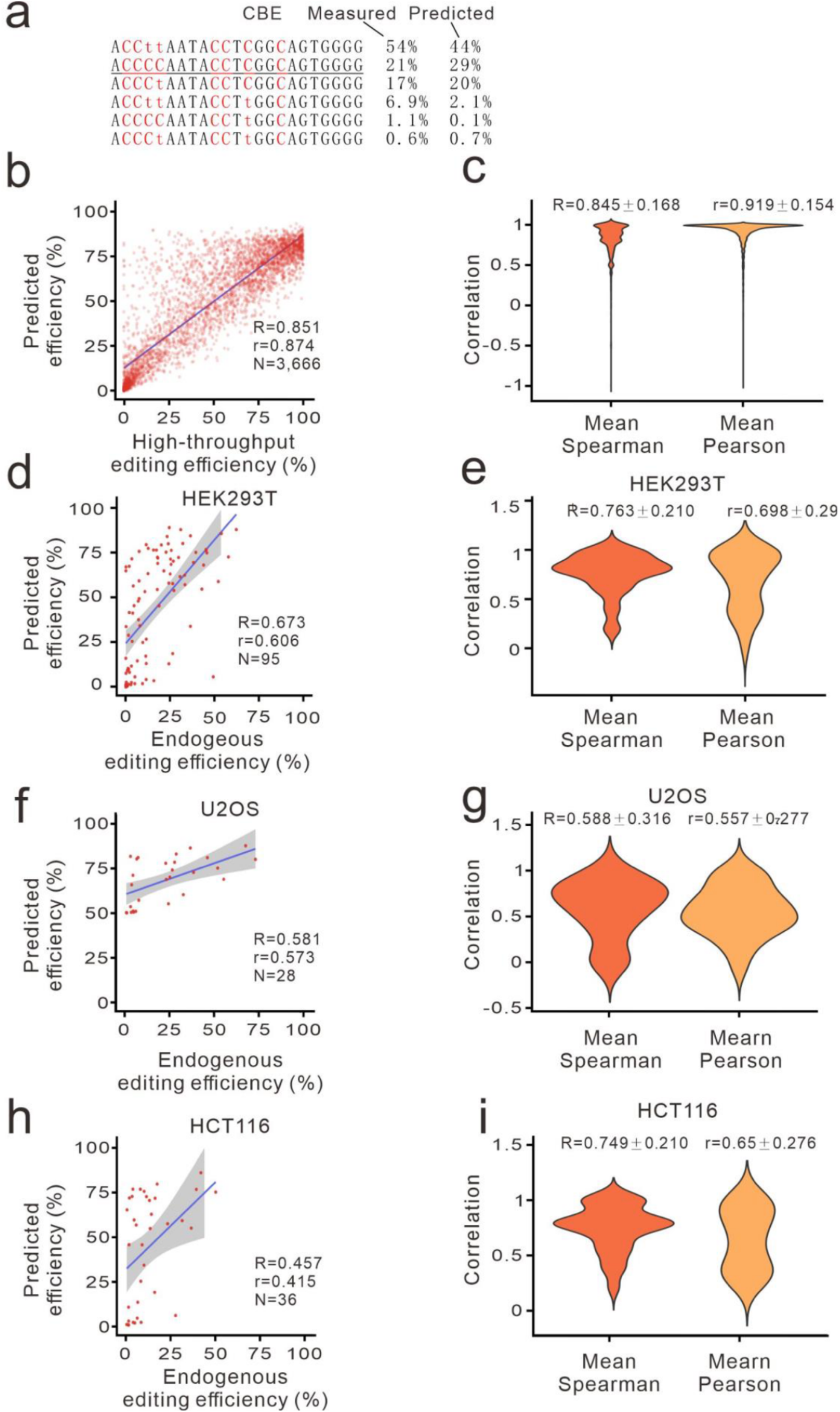
Evaluation of CBEdeepon for editing efficiency and outcome prediction. (**a**) Example of CBEdeepon prediction for a given target. Measured efficiencies are listed for comparison. Original target sequence is underlined. (**b**) Evaluation of CBEdeepon for conversion efficiency with testing dataset. (**c**) Evaluation of CBEdeepon for conversion outcome sequence frequencies with testing dataset. (**d, f, h**) Evaluation of CBEdeepon for conversion efficiency with endogenous dataset generated in HEK293T, U2OS and HCT116 cells, respectively. (**e, g, i**) Evaluation of CBEdeepon for conversion outcome sequence frequencies with endogenous dataset generated in HEK293T, U2OS and HCT116 cells, respectively.

Parallelly, we used the CBE editing datasets to train a shared embedding based deep learning model. The targets were randomly split to 68,244 (5,140,905 outcomes) for training, 1,393 (97,841 outcomes) for internal validation, and 3,666 (275,173 outcomes) held out for testing. The resulting model was named “CBEdeepon” which can predict editing outcome sequence frequencies (Fig. 4a). CBEdeepon achieved a Spearman correlation of 0.851 (r=0.874, MSE=0.027) for prediction of editing efficiency with testing dataset (Fig. 4b). To evaluate the performance for prediction of outcome frequency distribution, we tested 3,179 targets with at least 4 outcomes and achieved mean Spearman correlation of 0.845 (r=0.919, MSE=0.009, KL divergence=0.005, Fig. 4c). To test whether CBEdeepon works in other cell types, we used library to generate datasets in HeLa cells and obtained 56,529 valid target with 4,213,361 outcomes (reads number ≥100). We extracted 2,827 targets present in HaLa testing dataset as external testing dataset and achieved Spearman correlation of 0.864 (r=0.761, MSE=0.110) for efficiency prediction, and mean Spearman correlation of 0.788 for outcome distribution prediction (r=0.736, MSE=0.042, KL divergence=0.012, Supplementary Fig. 6a-b).

To evaluate the performance of CBEdeepon for endogenous targets, we collected endogenous editing data generated in HEK293T, U2OS and HCT116 cells from literature ^15^ and evaluated with our model, achieving mild correlation for efficiencies (Spearman correlation: 0.457 to 0.673) and high correlation for outcomes (mean Spearman correlation: 0.588-0.763, Fig.4 d-i). To evaluate the performance of the CBEdeepon in induced pluripotent stem cells (iPSCs), we generated dataset for 24 endogenous targets in human iPSCs. CBEdeepon achieved mild Spearman of 0.525 for efficiency prediction and mean Spearman correlation of 0.558 for outcome frequency prediction (Supplementary Fig. 6c-d).

We finally investigated the positional effects of target nucleotides on prediction. ABEdeepon achieved a good Spearman correlation (>0.5) from position 4 to 12, whereas CBEdeepon achieved a good Spearman correlation (>0.5) from position 3 to 18 (Supplementary Fig. 7a-b). Altogether, our results demonstrated that both ABEdeepon and CBEdeepon models perform well for prediction of conversion efficiency and outcome sequence frequencies.

Recently, the advent of three developed predictive model are also avaliable to predicting base editing outcomes, BE-Hive, DeepBaseEditor and BE-DICT, which are based on deep conditional autoregressive model, a two-hidden layer convolutional neuro network and a multi-head self-attention machanism seperately. The former two and ours have the characterize to seperately predict proportion of editing outcomes and overall editing efficiency, while the latter one gives the nature of predicting all editing outcomes containing overall efficiency directly. All of the previous models claimed that they can perform well on predicting, whereas there was no clear and explicit comparison of their ability to estimate editing outcomes on the criteria of internal predicting performance by each targeted sequence instead of external performance. Thus, we came up with a new benchmarking method among all above three model and ours based on two criteria: internally grouped predicting correlation and externally total predicting performance, and both of them were evaluated on all outcomes and edited outcomes(Fig. 5).

**Figure 5.**
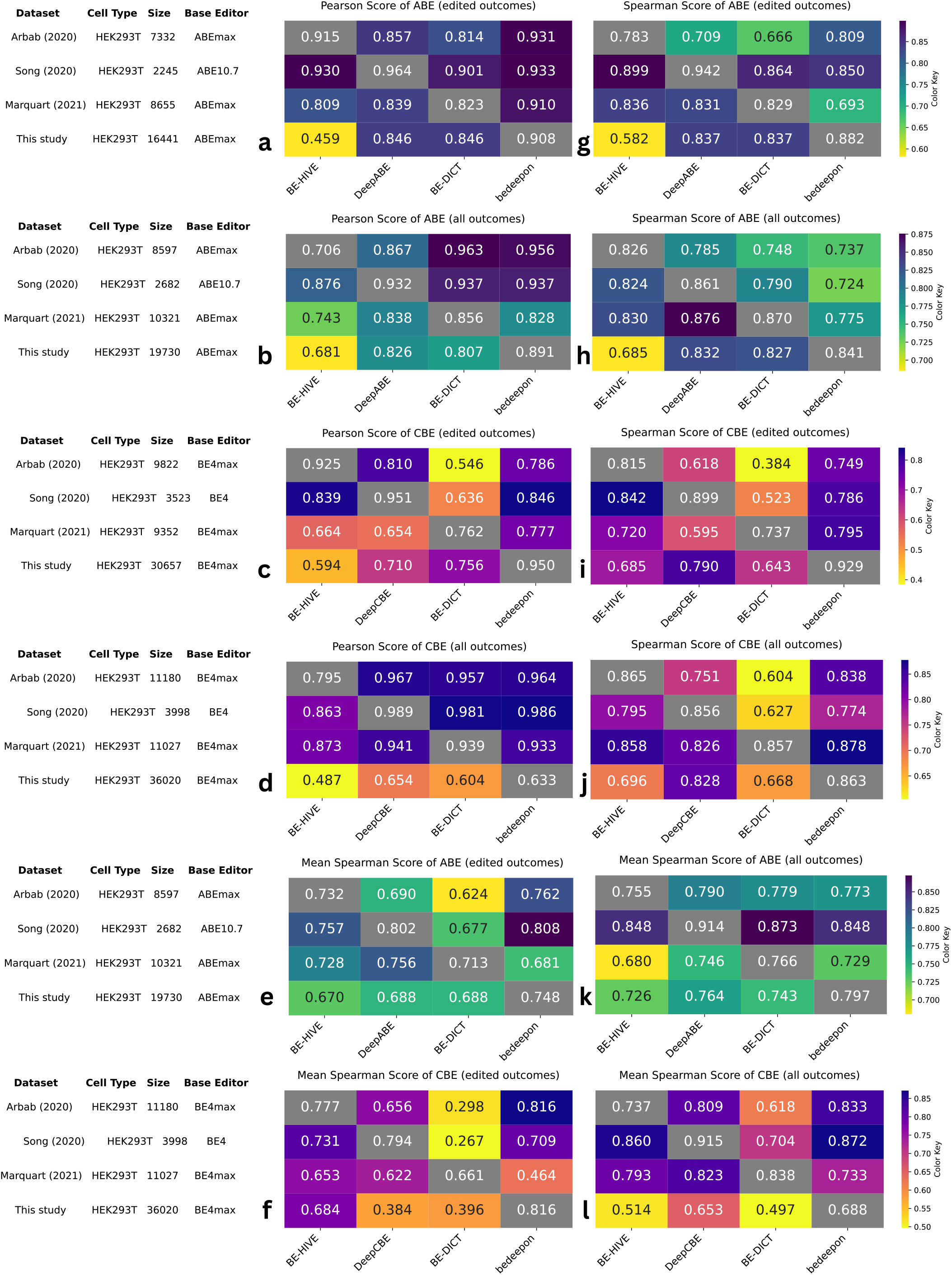
Benchmarking four model on two criterias of both internal and external coefficiency. Model benchmark evaluation is tested on 4 dataset from Arbab at el., Song at el., Marquart at el. and this study, the workflow is exhibit in Fig.1d. (**a-d,g-j**) Benchmarking performance with criteria one: calculating pearson and spearman correlation coefficient on overall predicted results; (**e,f,k,l**) criteria two: calculating correlation coefficient on mean spearman score of each group. The information of dataset reference, cell type, data size and the type of base editor are included in the left side.

To begin with, we compared our model with other three on the externally predicting ability to estimate all editing outcomes (with wild type), and we found our model perform the best as long as BE-Hive performed the worst on both ABE and CBE editing prediction(Fig. 5a,b). Additionally, the utility of predicting the proportion of edited-only outcomes (with no wild type) were also considered and our model outperform other three models without any doubt. Most importantly, rather than fluctuated Pearson and Spearman score, our bedeepon sustain a steady performance on four testing dataset, which manifested itself a outstanding generalizaion performance. Secondly, we put the newly internal criteria on these four models to calculate their mean spearman correlation coefficient, which was able to conceal a more reasonable and general ability of predicting. Consequently, a more solid conclusion for model performance were brought up with combining of the two criteria. In conclusion, our model still has the most outstanding and steady result while BE-DICT fluctuates the worst on both ABE and CBE by constrast.

## Discussion

Successful application of base editors requires a rigorous understanding of intended and unintended genome editing. Several groups have offered online tools or software for gRNA design, including iSTOP ^16^, beditor ^17^, BE-Designer ^18^ and BEable-GPS ^19^. These design rules conform to base editing requirements that the editable base falls within maximum activity window, but none of them can predict the editing activity and outcome frequencies. To solve this problem, we proposed shared embedding based deep learning models, ABEdeepon and CBEdeepon, to predict editing efficiencies and outcomes. Our model guided high performance for both prediction of base editing outcomes (mean R: 0.79-0.87 for high-throughput datasets) and efficiencies (R: 0.85-0.89 for high-throughput datasets).

During our manuscript revision, three groups developed tools named BE-Hive ^13^, DeepBaseEditor ^14^ and BE-DICT which achieved comparable performance to our models for prediction of editing efficiencies and outcomes recently. Consequently, we launched a series of innovative experiment to compare these four models extensively and found their models were all insteady. BE-Hive used carefully designed hand-crafted features to develop machine learning models for prediction of base editing outcomes and efficiency.

DeepBaseEditor followed the author’s previous method ^20^, extracting features using convolutional neural networks, which also achieved similar performance for prediction of base editing outcomes and efficiency ^14^. And BE-DICT developed a novel model with multi-head attention machanism to predict all outcomes which was edited or not. The intergration of all these models will enhance the prediction ability and facilitate selection of targets with high efficiency and desirable outcome sequences.

Overall, this shared embedding based deep learning models used here has a simple yet elegant structure which unifies on-target and off-target input in form. It can be seamlessly extended to other versions of base editors, such as SauriABEmax, SauriBE4max, SaKKH-BE3, BE4-CP, dCpf1-BE and eA3A-BE3 ^21-25^. Notably, it may also be used to generate models to predict editing outcomes for Cas9 and Cas12 nucleases.

## Methods

### Cell culture and transfection

HEK293T cells and HeLa cells (ATCC) were maintained in Dulbecco’s Modified Eagle Medium (DMEM) supplemented with 10% FBS (Gibco), while human iPSCs were cultured on Matrigel-coated plates (ESC qualified, BD Biosciences, San Diego, CA) using hESC mTeSR-1 cell culture medium (StemCell Technologies, Vancouver, Canada) at 37 °C and 5% CO2. All media contained 100 U/ml penicillin and 100 mg/ml streptomycin. For transfection, HEK293T cells and HeLa cells were plated into 6-well plates, DNA mixed with Lipofectamine 2000 (Life Technologies) in Opti-MEM according to the manufacturer’s instructions, while iPSCs using Lipofectamine 3000 (Life Technologies). Cells were tested negative for mycoplasma.

### Plasmid construction

SB transposon (pT2-SV40-BSD-ABEmax and pT2-SV40-BSD-BE4max) was constructed as follows: first, we replaced the Neo^R^ gene (AvrII-KpnI site) on pT2-SV40-Neo^R^ with BSD, resulting in pT2-SV40-BSD vector; second, backbone fragment of pT2-SV40-BSD was PCR-amplified with Gibson-SV40-F and Gibson-SV40-R, and ABEmax fragment was PCR-amplified from pCMV_ABEmax_P2A_GFP (Addgene#112101) with Gibson-ABE/BE4-F and Gibson-ABE/BE4-R, and BE4max fragment was PCR-amplified from pCMV_AncBE4max (Addgene#112094) with Gibson-ABE/BE4-F and Gibson-ABE/BE4-R; third, the backbone fragments were ligated with ABEmax and BE4max using Gibson Assembly (NEB), resulting in pT2-SV40-BSD-ABEmax and pT2-SV40-BSD-BE4max, respectively.

### Generation of cell lines expressing ABEmax or BE4max

HEK293T cells and HeLa cells were seeded at ∼40% confluency in a 6-well dish the day before transfection, 2 μg of SB transposon (pT2-SV40-BSD-ABEmax or pT2-SV40-BSD-BE4max) and 0.5 μg of pCMV-SB100x were transfected using 5 μl of Lipofectamine 2000 (Life Technologies). After 24 h, cells were selected with 10 μg/ml of blasticidin for 10 days. Single cells were sorted into 96-well plates for colony formation. Conversion efficiency was performed to screen cell clones with high levels of ABEmax and BE4max expression.

### The gRNA-target library construction

The on-target library design and construction were described previously^10^. Briefly, full-length oligonucleotides were PCR-amplified and cloned into Lentiviral vector by Gibson Assembly (NEB). The Gibson Assembly products were electroporated into MegaX DH10B^™^ T1^R^ Electrocomp^™^ Cells (Invitrogen) using a GenePulser (BioRad) and grown at 32 °C, 225 rpm for 16 h. The plasmid DNA was extracted from bacterial cells using Endotoxin-Free Plasmid Maxiprep (Qiagen).

### Lentivirus production

Lentivirus production was described previously^10^. Briefly, 12 μg of plasmid library, 9 μg of psPAX2, and 3 μg of pMD2.G (Addgene) were transfected into a 10-dish HEK293T cells with 60 μl of Lipofectamine 2000. Virus were harvested twice at 48 h and 72 h post-transfection. The virus was concentrated using PEG8000 (no. LV810A-1, SBI, Palo Alto, CA), dissolved in PBS and stored at −80 °C.

### Screening experiments in HEK293T and HeLa cells

HEK293T or HeLa cells expressing ABEmax or BE4max were plated into 15 cm dish at

∼30% confluence. After 24 h, cells were infected with gRNA library with at least 1000-fold coverage of each gRNAs. After 24 h, the cells were cultured in the media supplemented with 2 μg/ml of puromycin for 5 days. Cells were harvested and the genomic DNA was isolated using Blood & Cell Culture DNA Kits (Qiagen). The integrated region containing the gRNA coding sequences and target sequences were PCR-amplified using primers Deep-seq-library-F/R with Q5 High-Fidelity 2X Master Mix (NEB). We performed 60-70 PCR reactions using 10 μg of genomic DNA as template per reaction for deep sequencing analysis. The PCR conditions: 98 °C for 2 min, 25 cycles of 98 °C for 7 s, 67 °C for 15 s and 72 °C for 10 s, and the final extension, 72 °C for 2 min. The PCR products were mixed and purified using Gel Extraction Kit (Qiagen). The purified products were sequenced on Illumina HiSeq X by 150-bp paired-end sequencing.

### Data analysis

FASTQ raw sequencing reads were processed to identify gRNA editing activity and editing outcomes. The nucleotides in a read with quality score < 10 was masked with a character “N”. Due to the integrated design strategy, we first separated a read to designed gRNA region, scaffold region, and target region to extract the corresponding sequence. The designed gRNA was then aligned to the reference gRNA library to mark the reads. The target sequence was compared to the designed gRNA to mark all types of conversion (i.e., canonical A-G, C-T conversions and non-canonical A-C/T, C-A/G, G-A/T, T-A/C conversions at each position). We screened out gRNAs with a total valid reading of less than 100. Then, the efficiency for a specific conversion type at a position can be calculated by the following formula:

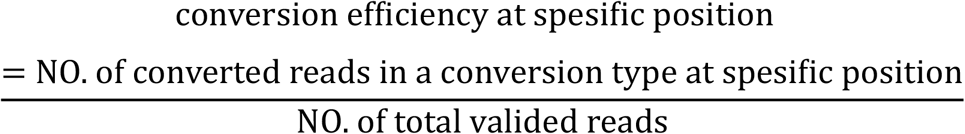

However, the conversion efficiency was very low for non-canonical conversions (mean efficiency < 0.005). Thus, we only consider the canonical conversions for the outcome frequency distribution analysis. Theoretically, the canonical editing combinations of 20 bases is at most 2^20^. However, we found that positions 1-2 and 18-20 always contain only one conversion. If there exist canonical bases, positions 3-17 may have multiple conversions at the same time. Therefore, we programmed this conversion scheme to obtain all possible editing outcomes and assign true editing frequencies to the outcomes that exist in the sequencing data and assign 0 frequencies to the outcomes not found in the sequencing data. Then, the editing frequency of a gRNA-outcome can be described as:

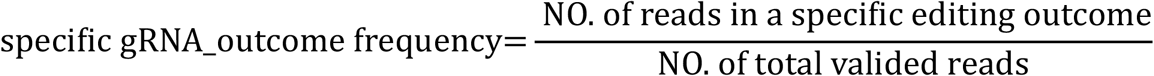

Note that the non-converted targets were also considered as an editing outcome. So, the editing efficiency for a target can be simply calculated by the following formula:

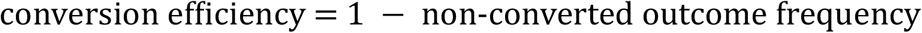

### Encoding

Drawing on concepts from the field of natural language processing (NLP), nucleotides A, C, G, and T can be regarded as words in a DNA sequences. Therefore, we can widely use algorithms in the NLP field to solve prediction tasks in the CRISPR field, especially the use of embedding algorithms to get the continuous representation of discrete nucleotide sequences ^26^. Unlike the common efficiency prediction that only needs to input one single sequence for regression models, in this research, both the gRNA-outcome pairs (for BEdeepon) and gRNA-target pairs (for BEdeepoff) has two different sequences as inputs. For gRNA-outcome pairs, there are four words in the index vocabulary (i.e., A, C, G, and T). So, the vocabulary can be described as: So, an input sequence can be described as: where *i* ∈ 1,2denotes the *i*-th sequence in an gRNA-outcome pair or an gRNA-target pair, *x*_i*t*_ is the *t*-th element of the *i*-th sequence, *T* is the sequence length. For example, for a gRNA-outcome pair GTGGAACATCCACTTGACCTAGG (seq1, gRNA + NGG) and GTGGAGCGTCCACTTGACCTAGG (seq2, one outcome) can be encoded as:

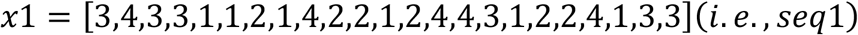

and

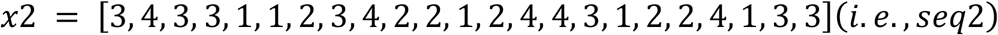

respectively.

### Shared embedding

Inspired by the algorithms in the recommender system ^27^ and click-through rate (CTR) ^28^ prediction modeling, both the generalization capacity and training speed will benefit from the sharing of the same embedding matrix instead of training independent embedding matrices for each input. In this research, a discrete nucleotide encoding *x*_*it*_ is projected to the dense real-valued space ***E***_***i***_ ∈ ℝ^*T*×*m*^ (*m* is a hyperparameter corresponds to the embedding dimension) to get the embedding vector ***e***(*x*_*it*_). Then a final embedding matrix ***E*** is needed to get the combined information from those two embedding matrices by:

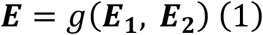

where *g* can be sum, mean, or even a simple concatenate function. However, the sum or mean function is more suitable because it can reduce the redundant features in ***E***_**1**_ and ***E***_**2**_. We choose the sum function here for simplicity.

### Feature extraction and model prediction

Long-short-term memory network (LSTM) and gated recurrent unit network (GRU) are a type of recurrent neural networks (RNN) algorithms used to address the vanishing gradient problem in modelling time-dependent and sequential data tasks ^29^. Usually, a bidirectional manner was used to capture the information from the forward and backward directions of a sequence, which is biLSTM or biGRU. Our work and others’ work have shown that, as an important component, biLSTM can be used alone or with convolutional neural network (CNN) to achieve good performances in various regression and classification tasks involving biological sequences ^10, 30-32^. Here, we tried biLSTM, biGRU, and the newly proposed transformer structure ^33^, and found biLSTM had the fastest convergence speed. The input and output of biLSTM can be described by the following equations:

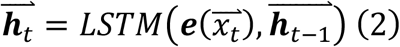

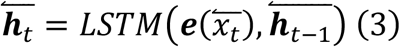

Thus, the output context vectors of biLSTM are 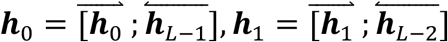 etc. Thus, we can concatenate the forward and backward hidden state as***H*** = ***h***_**0**_, ***h***_**1**_,…, ***h***_***L*−1**_,which contains the bidirectional information in the shared embedding feature matrix. Before the fully connected layers, we tried different input features based on the trade-off of the convergence speed and the performance of the model. The aforementioned features are last hidden unit, max pooling operation on ***H***, and average pooling on ***H***. The equations are as following:

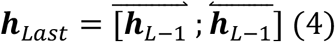

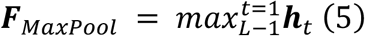

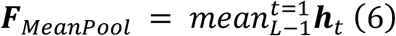

We observed that the last hidden state is the only need to obtain an optimal performance, i.e.,

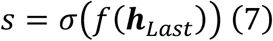

where, *f* is fully connected layers, *σ* is the *LeakyReLU* activation function and *o* is the output score for a specific gRNA-outcome pair.

It should be noted that the gRNA + NGG in a set of gRNA-outcome pairs (a gRNA batch with *K* samples) are all the same, and the outcome sequences are converted from the same target, so the output scores of a gRNA-batch can be denoted as ***s*** = [*s*_1_, *s*_2_, …, *s*_*k*_]. Then, a *softmax* activation function can be applied to ***s*** to get the predicted frequency distribution with a sum of 1 for the gRNA-outcome pair (Eq. 8, 9).

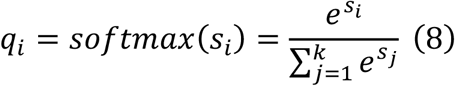

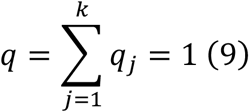

We have described in the **Data Analysis** section, that the conversion efficiency of a gRNA can be calculated by the formula:

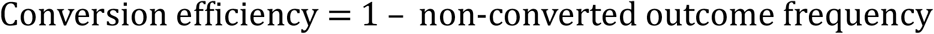

If we let *q*_0_ be non-converted outcome frequency in ***q***. Then, the predicted efficiency will be 1 – *q*_0_.

### Combined weighted loss function

For the on-target models, a calculated true gRNA-outcome frequency distribution can be denoted as ***p*** = [*p*_1_, *p*_2_, …, *p*_*K*_]. So, it’s naturally to apply Kullback–Leibler (KL) divergence loss function to minimize the difference between the predicted frequency distribution ***q*** and the true frequency distribution ***p***. The standard KL divergence loss function *D*_*KL*_(***p***||***q***) is defined as:

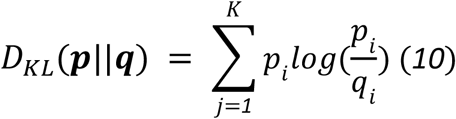

Basically, we wanted that a gRNA-outcome pair with a large number of reads in a gRNA batch has a more accurate prediction value. The weight of a sample loss was re-assigned depending on its corresponding read counts *w*_*i*_. The modified loss function of Eq.10 is:

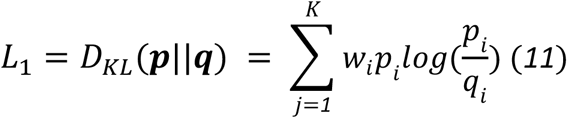

A number of studies ^34^ have shown that multi-task learning architecture can significantly improves the stability and generalization capacity of the model. In addition to KL divergence loss (which measures the difference between the two probability distributions as a whole), the mean squared error (MSE) loss function also helps to minimize the difference between each of the *pi, qi* pairs individually. So, we adopted the MSE loss in a weighted manner:

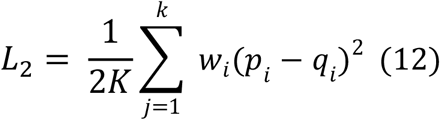

Then, the final loss function can be denoted as follows in Eq.13:

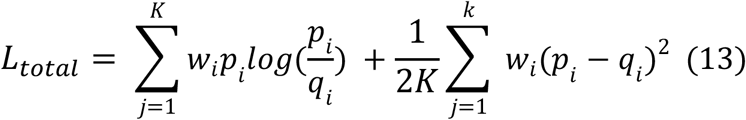

The loss function for the off-target model is simply the ordinary MSE loss.

### Training setting

The on-target datasets were randomly split into two parts with ratio 9 : 1, the former for training, and the latter for holdout testing. To made a more stable prediction, we not only adapted cross-validation training for this dataset but also used more external datasets to test the generalization capacity of the models. The training sample size of our on-target datasets are 29,604 on ABE and 48,270 on CBE, which are concatenated with other three datasets: 8261 on ABE and 8029 on CBE (Arbab at el.), 10,209 on ABE and 9,743 on CBE (Song at el.), 6,891 on ABE and 7,859 on CBE (Marquart at el.). Then the stability of model performance was estimated by a 5-fold shuffled validation together with the external integrated and endogenous datasets. The ABEdeepon and CBEdeepon models share the following hyperparameters: embedding dimension, 128; BiLSTM hidden unites, 256; BiLSTM hidden layers, 2; dropout rate, 0.3; fully connected layers,1 (2 * 2 * 256 → 1). For the Adam optimizer, it was used with a customized learning rate decay strategy that gradually reduce the learning rate from 0.001, 0.0001, 0.00005 to 0.00001.

### Testing Configuration and criterias of benchmark

In order to solidify our conclusion of our model’s outstanding performance on estimating, we collected and integrated datasets from three previously published project: sample 1,265 on ABE and 1,358 on CBE (Arbab at el.), 437 on ABE and 475 on CBE (Song at el.), 1,667 on ABE and 1675 on CBE (Marquart at el.) and our split testing dataset, including sample 3,289 on ABE and 5,363 on CBE. In summary, an aggregated dataset with total sample 6,658 and 8,871 on ABE and CBE are adapted as “ground-truth” to benchmark individually.

In progress, we chose both “bystander” and “overall efficiency” models from BE-hive and DeepBaseEditor and “bystander” module in BE-DICT to predict on both edited outcomes (called proportion) and overall outcomes (called frequency). The way to calculating both proportion is aligned with 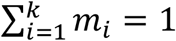, where *m* means edited outcomes; and frequency is aligned with 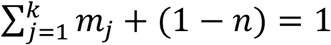, where *n* represents overall editing efficiency.

Considering the base editing happens on each targeted sequence, except compare the overall external correlation coefficient, a better evaluating criteria turns out with calculating mean spearman correlation coefficient within each group, and also the variance (Supplementary Table.14, Supplementary Figure.12-13).

### Tools used in the study

Bwa-mem2 was used to identify and align the designed gRNA^35^. pytorch==2.4.0 torchvision==0.19.0, torchaudio==2.4.0, pytorch-cuda=11.8 ^36^ was used for building deep learning models.

## Code availability

We provide the source code for bedeepon and the custom Python scripts used to train and evaluate the models and benchmarks available on GitHub at https://github.com/martina-yu/bedeepon and preprocessing procedure at https://github.com/martina-yu/PreprocessingBedeepon. The web for bedeepon in predicting both efficiency and proportion on the DNA sequence is published at http://www.deephf.com/#/bedeep/bedeepon.

## Supporting information

Supplemental figures and Tables are avaliable.

## Acknowledgments

This work was supported by grants from the National Natural Science Foundation of China (81870199, 81630087), the Foundation for Innovative Research Group of the National Natural Science Foundation of China (31521003), Science and Technology Research Program of Shanghai (Grant Number 19DZ2282100).

This pre-print is available under a Creative Commons License (Attribution-NonCommercial-NoDerivs 4.0 International), CC BY-NC-ND 4.0, as described at http://creativecommons.org/licenses/by-nc-nd/4.0/

## Notes

### Competing Interest Statement

The authors have declared no competing interest.

### Summary of Updates

Our revision has re-analyzed our raw data and brought up a new result initially. Then we have concatenated three previously published training dataset with ours to guide a more steady and generalized performance. Next, We came up with a novel criteria of benchmark and conducted a series of experiments to manifest itself within an outstanding integrating characterized performance. In conclusion, we recorded our innovative construction of model and a solid evaluation on it.

