## Supplementary materials for "BEdeepon: an in silico tool for prediction of base editor efficiencies and outcomes"

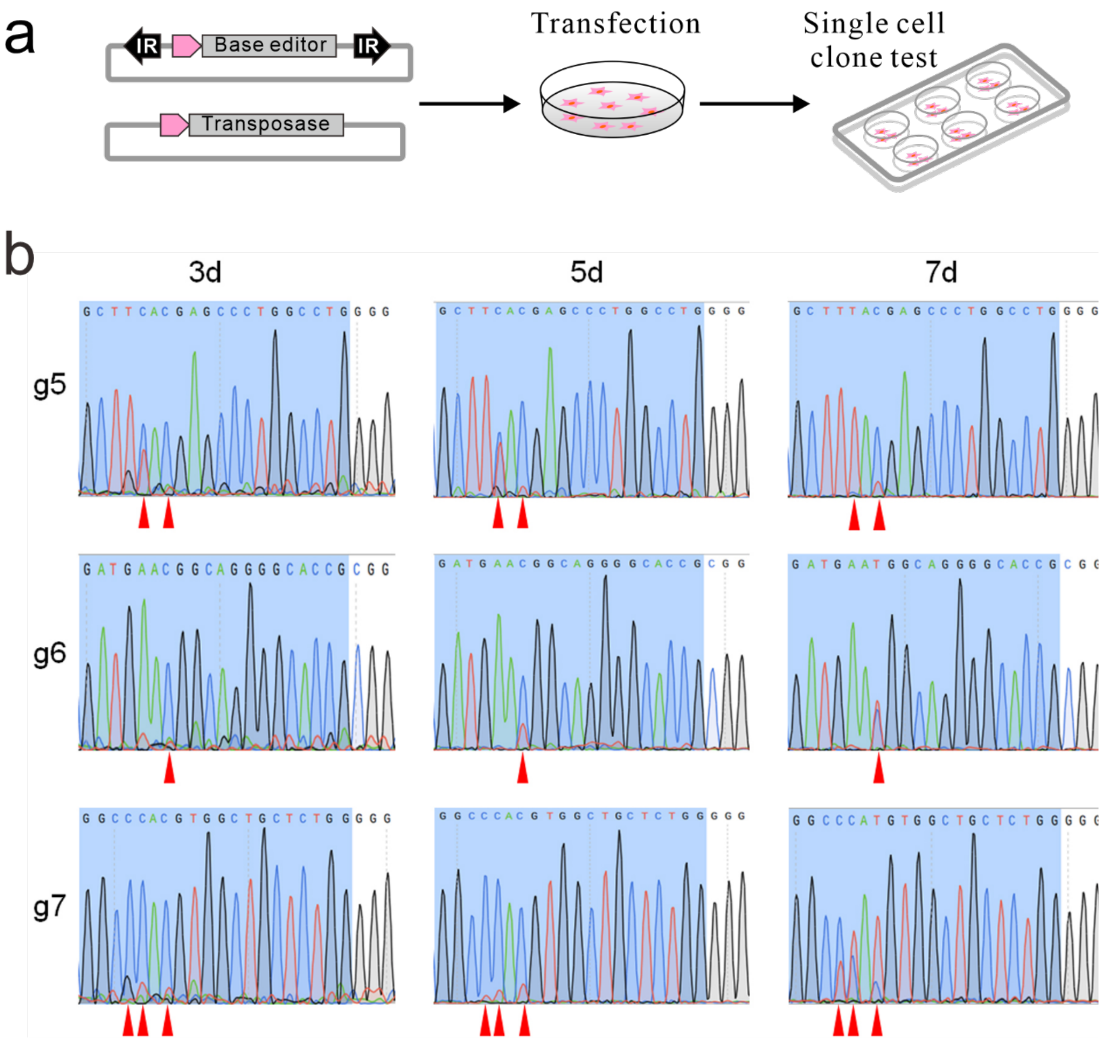


**Supplementary Figure 1. Generation of single cell-derived clones expressing base editors.**

(**a**) Schematic diagram of cell clone generation. The Sleeping Beauty transposon system, carrying base editors with transposase vector, is transfected into HEK293T cells followed by blasticidin selection. Five days after transfection, single cells are isolated and seeded into a new plate for colony formation. (**b**) Analysis of ABE base editing. Conversion efficiency at integrated sites was measured at days 3, 5 and 7 after gRNA-target paired lentivirus infection. The red arrow indicates the base of the conversion.

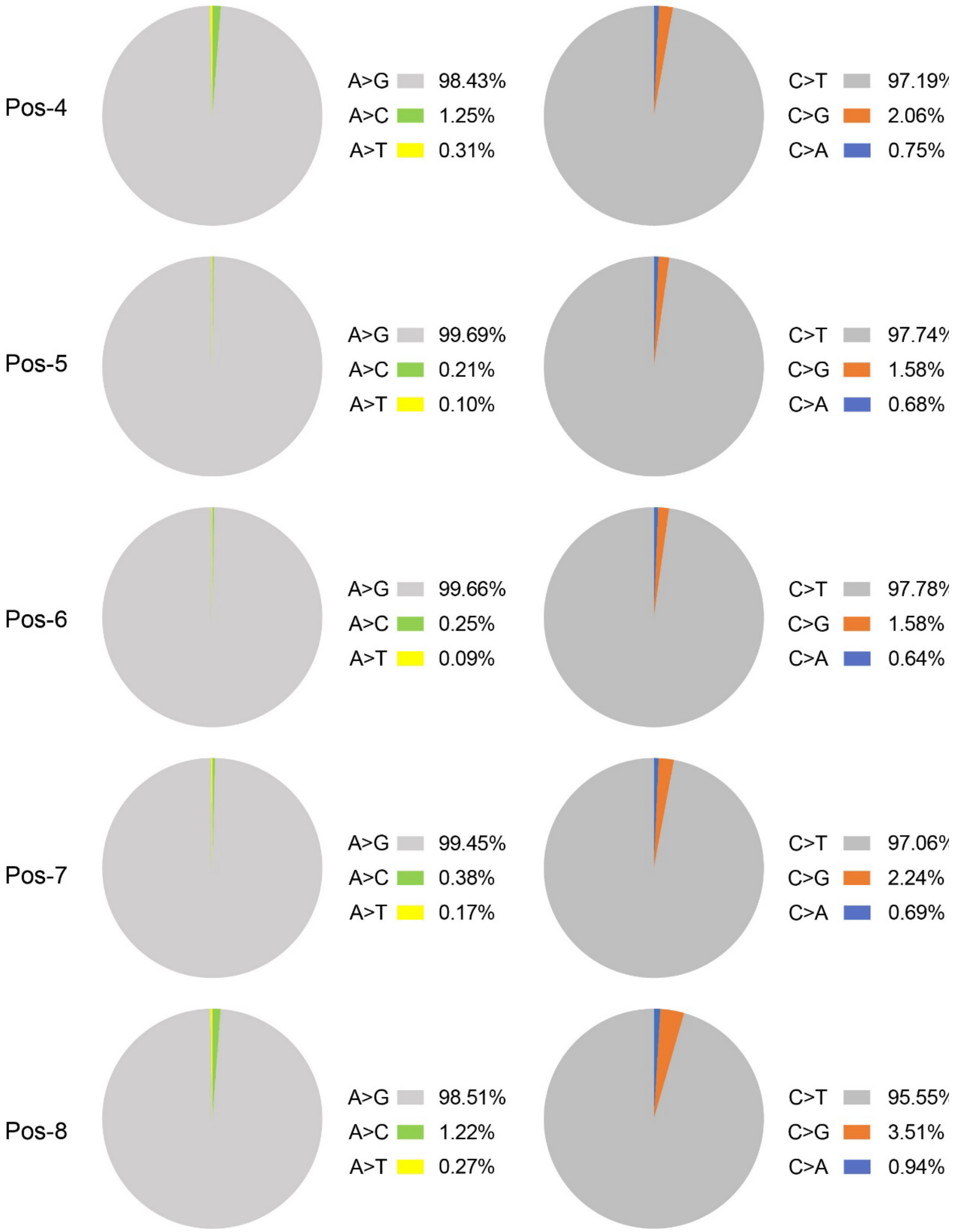


**Supplementary Figure 2. The product purity of nucleotide conversion for ABE (left) and CBE (right).**
The target nucleotide position is shown on the left.

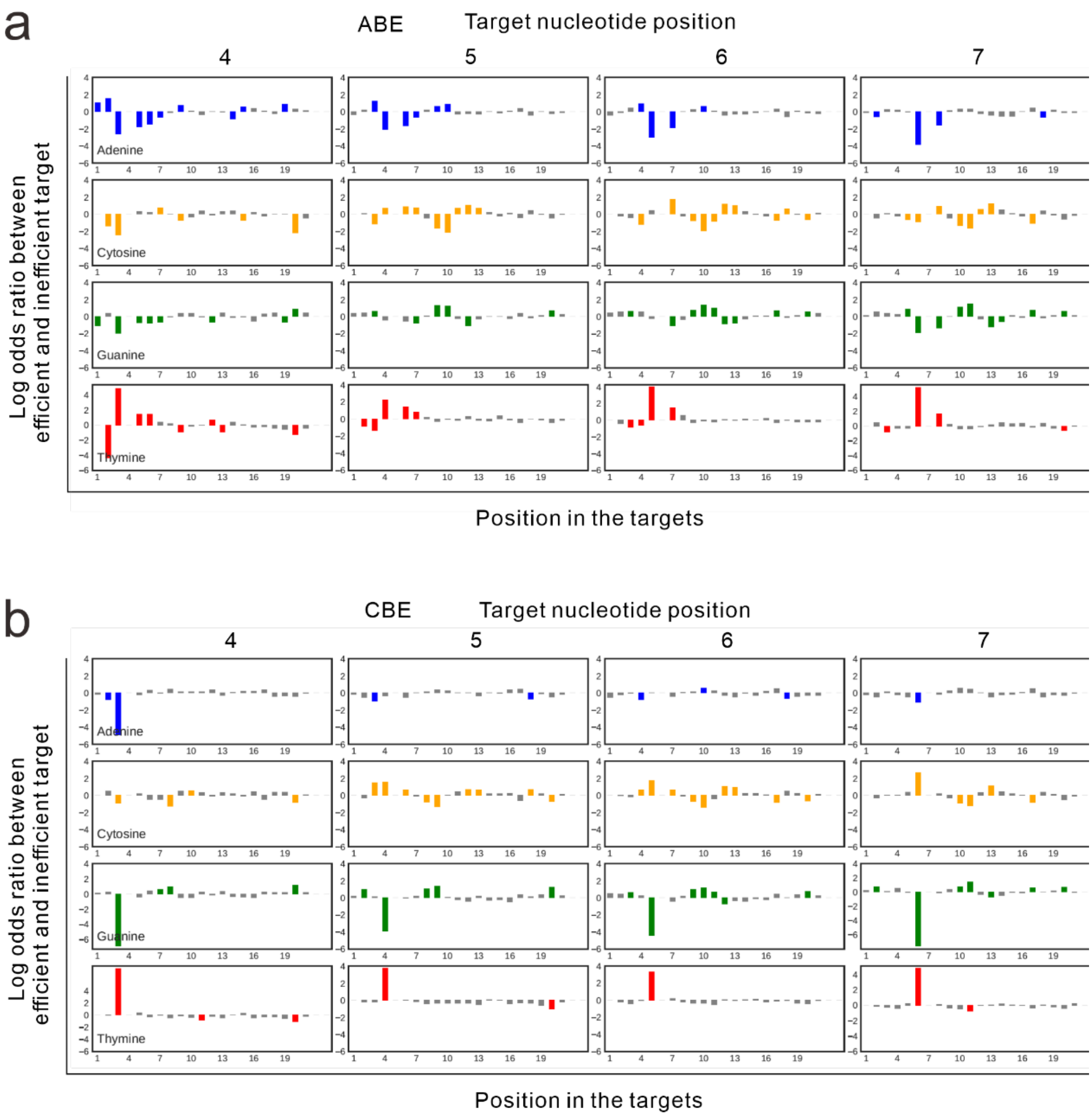


**Supplementary Figure 3. Sequence preferences at each position.** Sequence preferences at each position in efficient (top 20%) vs. inefficient (bottom 20%) targets for ABE (**a**) and CBE (**b**). The target nucleotide position is shown on the top. Nucleotide position on the targets is shown below. The log odds ratios of nucleotide frequencies between efficient and inefficient target sequences are represented on the y axis.

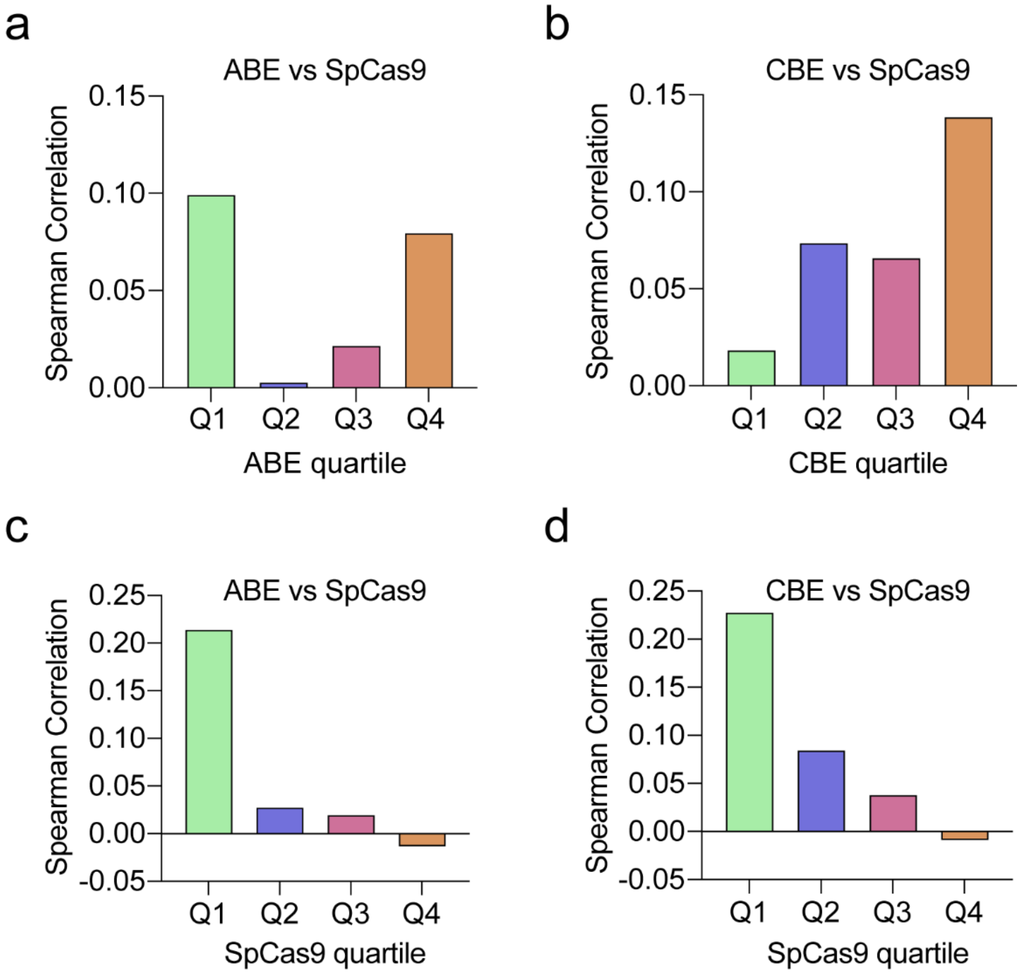


**Supplementary Figure 4. Spearman correlation of editing efficiency between SpCas9 and base editors.**

(**a**) Spearman correlation of editing efficiency between SpCas9 and ABE for each quartile of ABE editing efficiency. (**b**) Spearman correlation of editing efficiency between SpCas9 and CBE for each quartile of CBE editing efficiency. (**c**) Spearman correlation of editing efficiency between SpCas9 and ABE for each quartile of SpCas9 editing efficiency. (**d**) Spearman correlation of editing efficiency between SpCas9 and CBE for each quartile of SpCas9 editing efficiency.

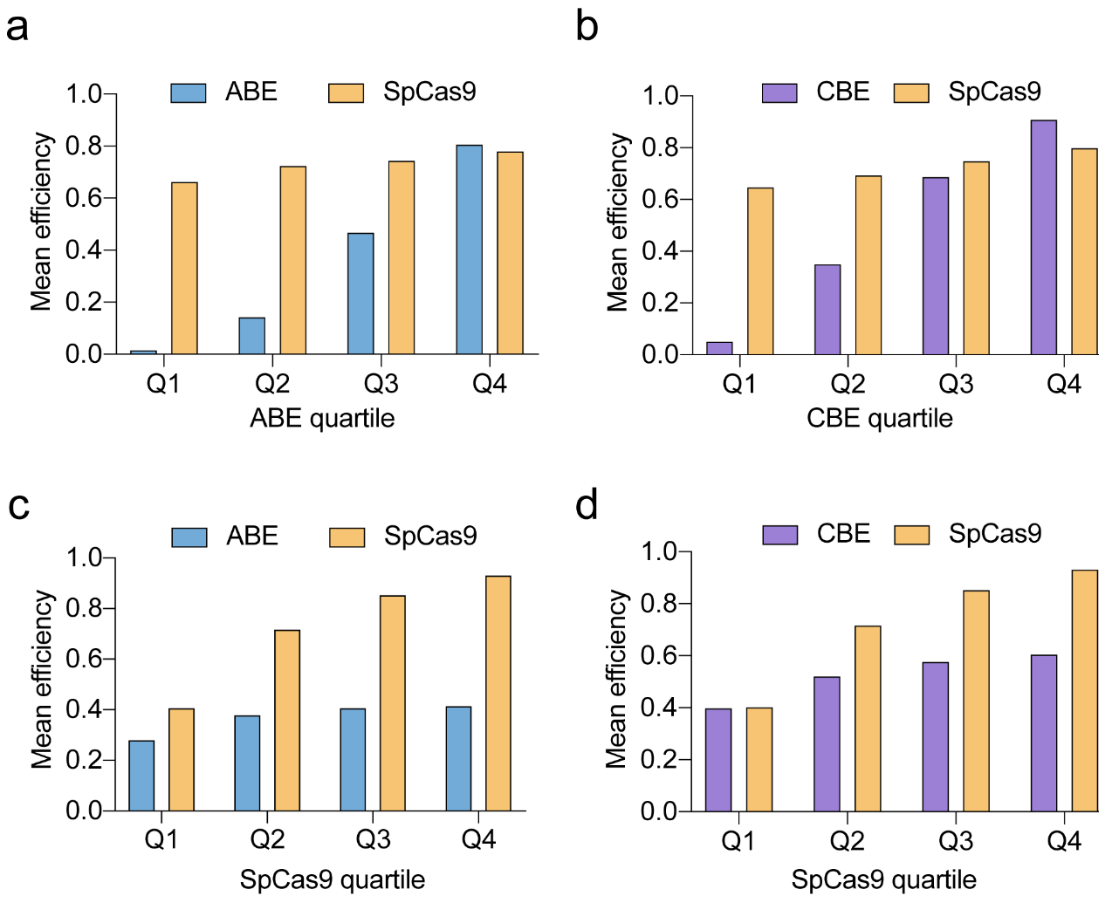


**Supplementary Figure 5. Mean editing efficiency comparison between SpCas9 and base editors.**

(**a**) Mean editing efficiency comparison between SpCas9 and ABE for each quartile of ABE editing efficiency. (**b**) Mean editing efficiency comparison between SpCas9 and CBE for each quartile of CBE editing efficiency. (**c**) Mean editing efficiency comparison between SpCas9 and ABE for each quartile of SpCas9 editing efficiency. (**d**) Mean editing efficiency comparison between SpCas9 and CBE for each quartile of SpCas9 editing efficiency.

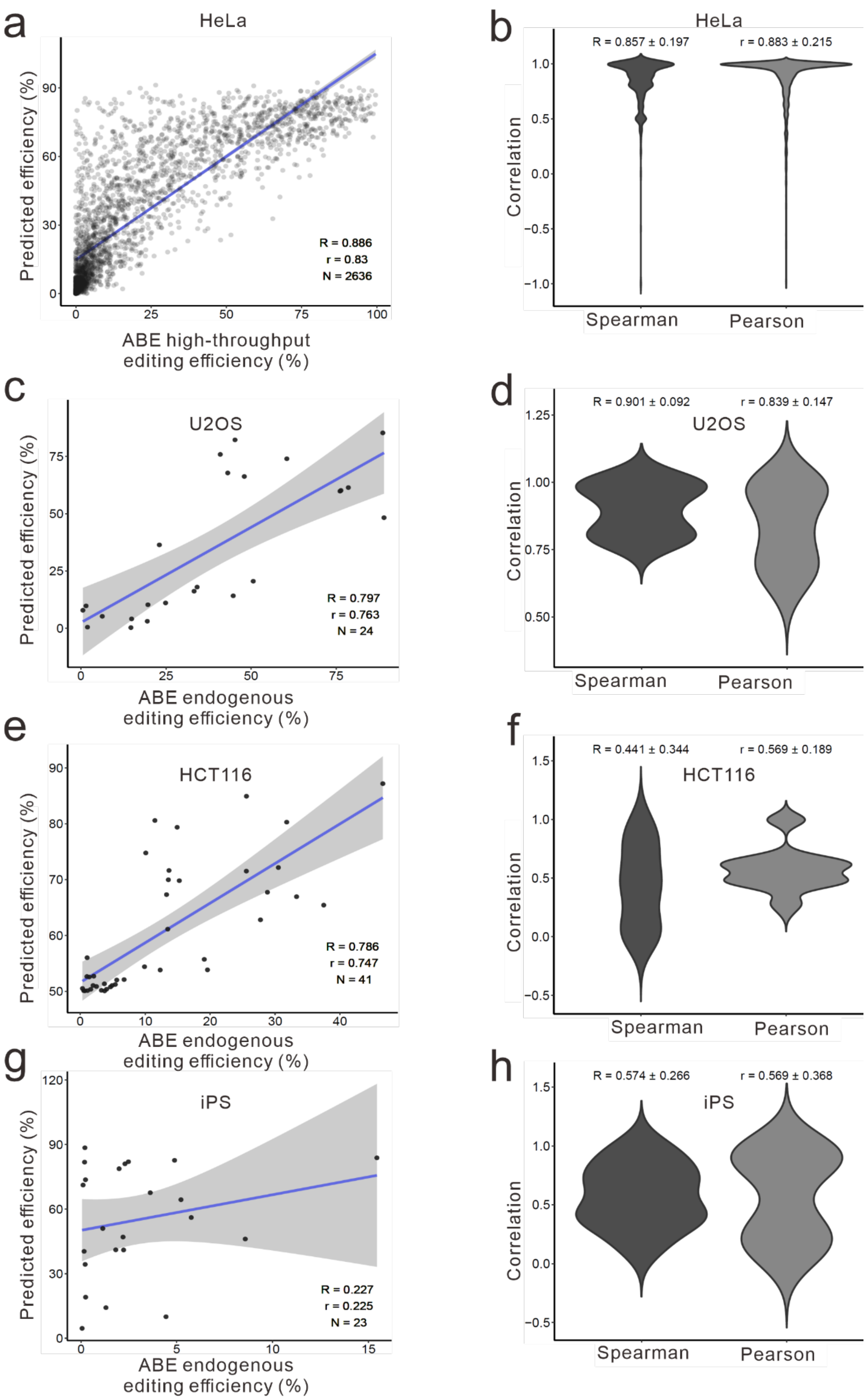
a
HeLa
Predicted efficiency (%)
90
60
30
0
R = 0.886
r = 0.83
N = 2636
0
25
50
75
100
ABE high-throughput
editing efficiency (%)
b
HeLa
R = 0.857 ± 0.197
r = 0.883 ± 0.215
Correlation
1.0
0.5
0.0
−0.5
−1.0
Spearman
Pearson
c
U2OS
Predicted efficiency (%)
75
50
25
0
R = 0.797
r = 0.763
N = 24
0
25
50
75
ABE endogenous
editing efficiency (%)
d
R = 0.901 ± 0.092
r = 0.839 ± 0.147
U2OS
Correlation
1.25
1.00
0.75
0.50
Spearman
Pearson
e
HCT116
Predicted efficiency (%)
90
80
70
60
50
R = 0.786
r = 0.747
N = 41
0
10
20
30
40
ABE endogenous
editing efficiency (%)
f
R = 0.441 ± 0.344
r = 0.569 ± 0.189
HCT116
Correlation
1.5
1.0
0.5
0.0
−0.5
Spearman
Pearson
g
iPS
Predicted efficiency (%)
120
90
60
30
0
R = 0.227
r = 0.225
N = 23
0
5
10
15
ABE endogenous
editing efficiency (%)
h
R = 0.574 ± 0.266
r = 0.569 ± 0.368
iPS
Correlation
1.5
1.0
0.5
0.0
−0.5
Spearman
Pearson

**Supplementary Figure 6. Evaluation of ABEdeepon for conversion efficiency and outcome frequency prediction.**

(**a**) Evaluation of ABEdeepon for conversion efficiency prediction at integrated targets in HeLa cells. (**b**) Evaluation of ABEdeepon for prediction of conversion outcome sequence frequencies at integrated targets in HeLa cells. (**c**) Evaluation of ABEdeepon for conversion efficiency prediction at endogenous targets in U2OS cells. (**d**) Evaluation of ABEdeepon for prediction of conversion outcome sequence frequencies at endogenous targets in U2OS cells. (**e**) Evaluation of ABEdeepon for conversion efficiency prediction at endogenous targets in HCT116 cells. (**f**) Evaluation of ABEdeepon for prediction of conversion outcome sequence frequencies at endogenous targets in HCT116 cells. (**g**) Evaluation of ABEdeepon for conversion efficiency prediction at endogenous targets in iPS cells. (**h**) Evaluation of ABEdeepon for prediction of conversion outcome sequence frequencies at endogenous targets in iPS cells.

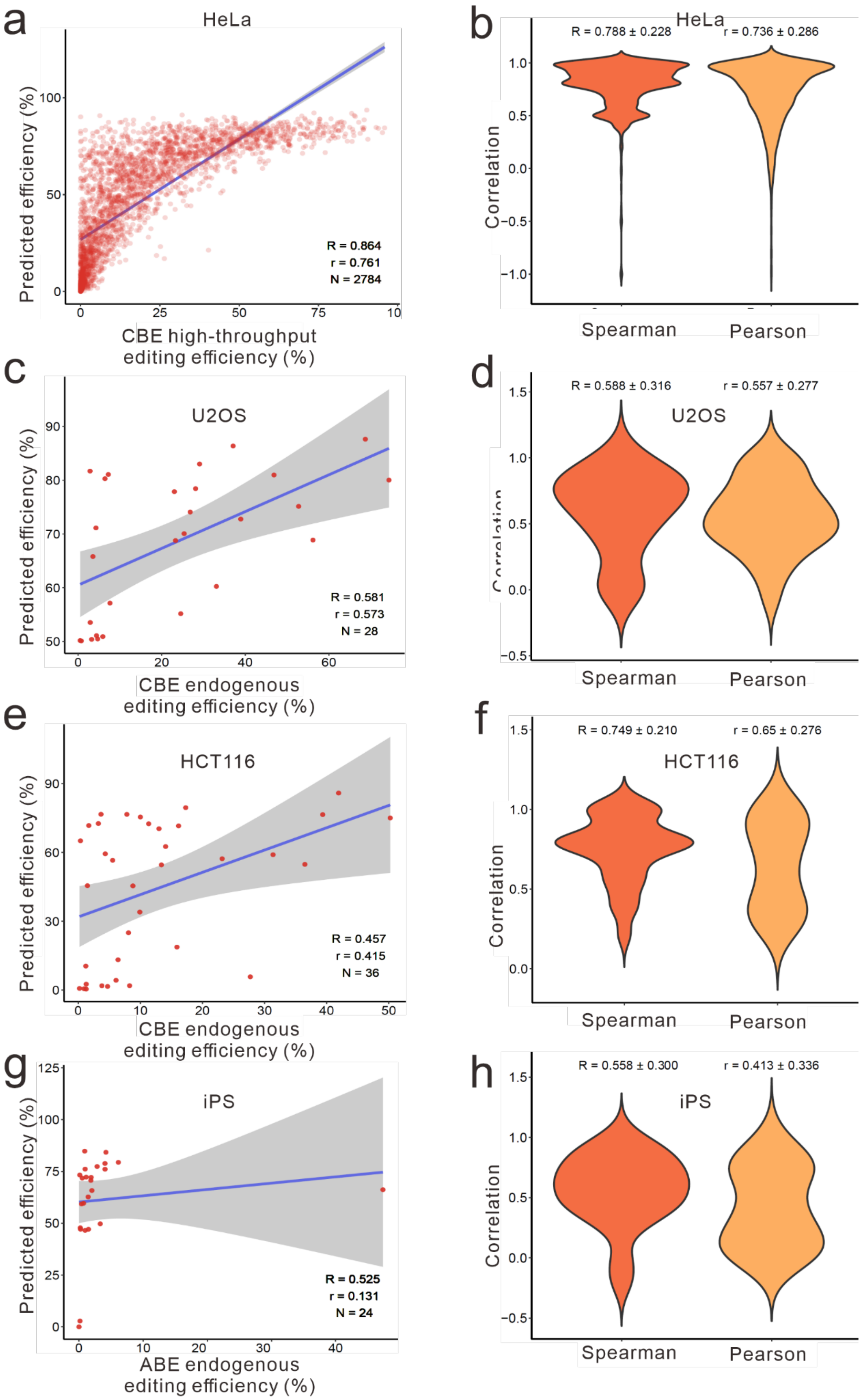


**Supplementary Figure 7. Evaluation of CBEdeepon for conversion**

**efficiency and outcome sequence prediction.**
(**a**) Evaluation of CBEdeepon for conversion efficiency prediction at integrated targets in HeLa cells. (**b**) Evaluation of CBEdeepon for prediction of conversion outcome sequence frequencies at integrated targets in HeLa cells. (**c**) Evaluation of CBEdeepon for conversion efficiency prediction at endogenous targets in U2OS cells. (**d**) Evaluation of CBEdeepon for prediction of conversion outcome sequence frequencies at endogenous targets in U2OS cells. (**e**) Evaluation of CBEdeepon for conversion efficiency prediction at endogenous targets in HCT116 cells. (**f**) Evaluation of CBEdeepon for prediction of conversion outcome sequence frequencies at endogenous targets in HCT116 cells. (**g**) Evaluation of CBEdeepon for conversion efficiency prediction at endogenous targets in iPS cells. (**h**) Evaluation of CBEdeepon for prediction of conversion outcome sequence frequencies at endogenous targets in iPS cells.

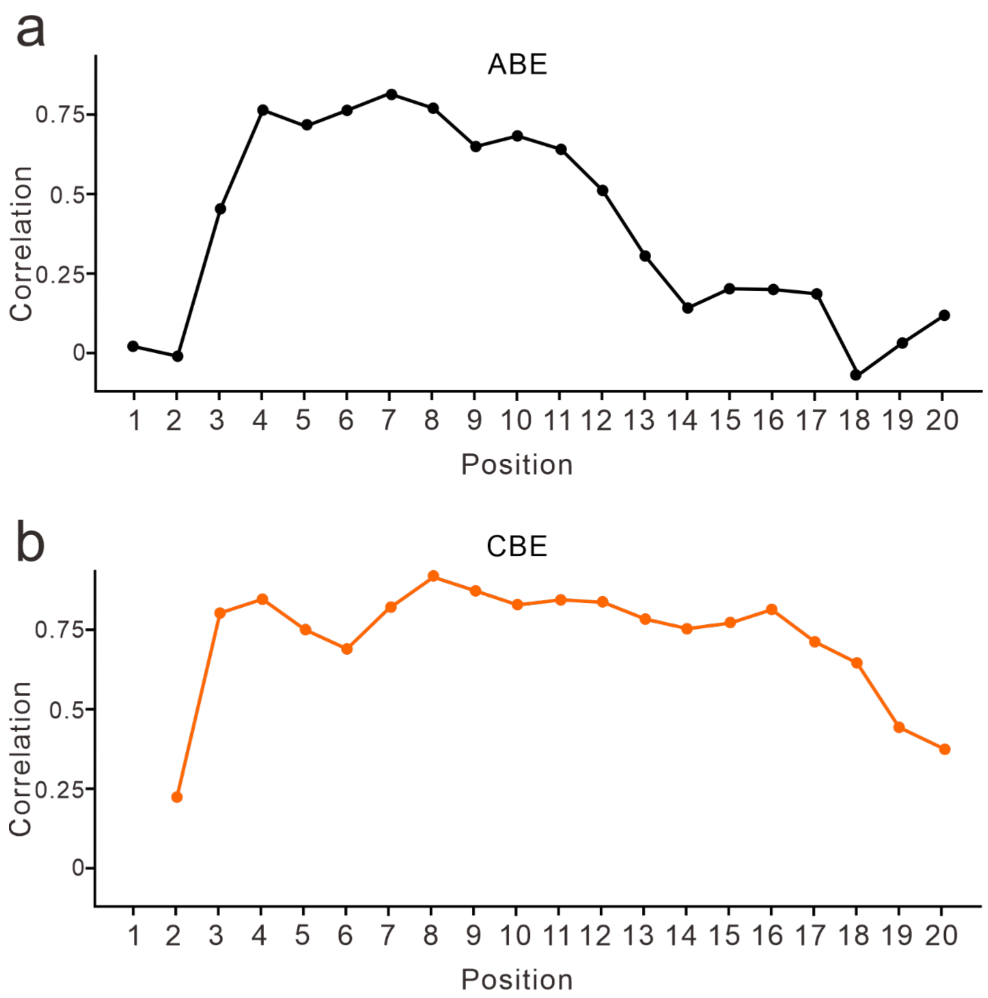


**Supplementary Figure 8. Positional effects on prediction performance.**
(**a**) Spearman correlation of ABEdeepon at different position of targets. (**b**) Spearman correlation of CBEdeepon at different position of targets.

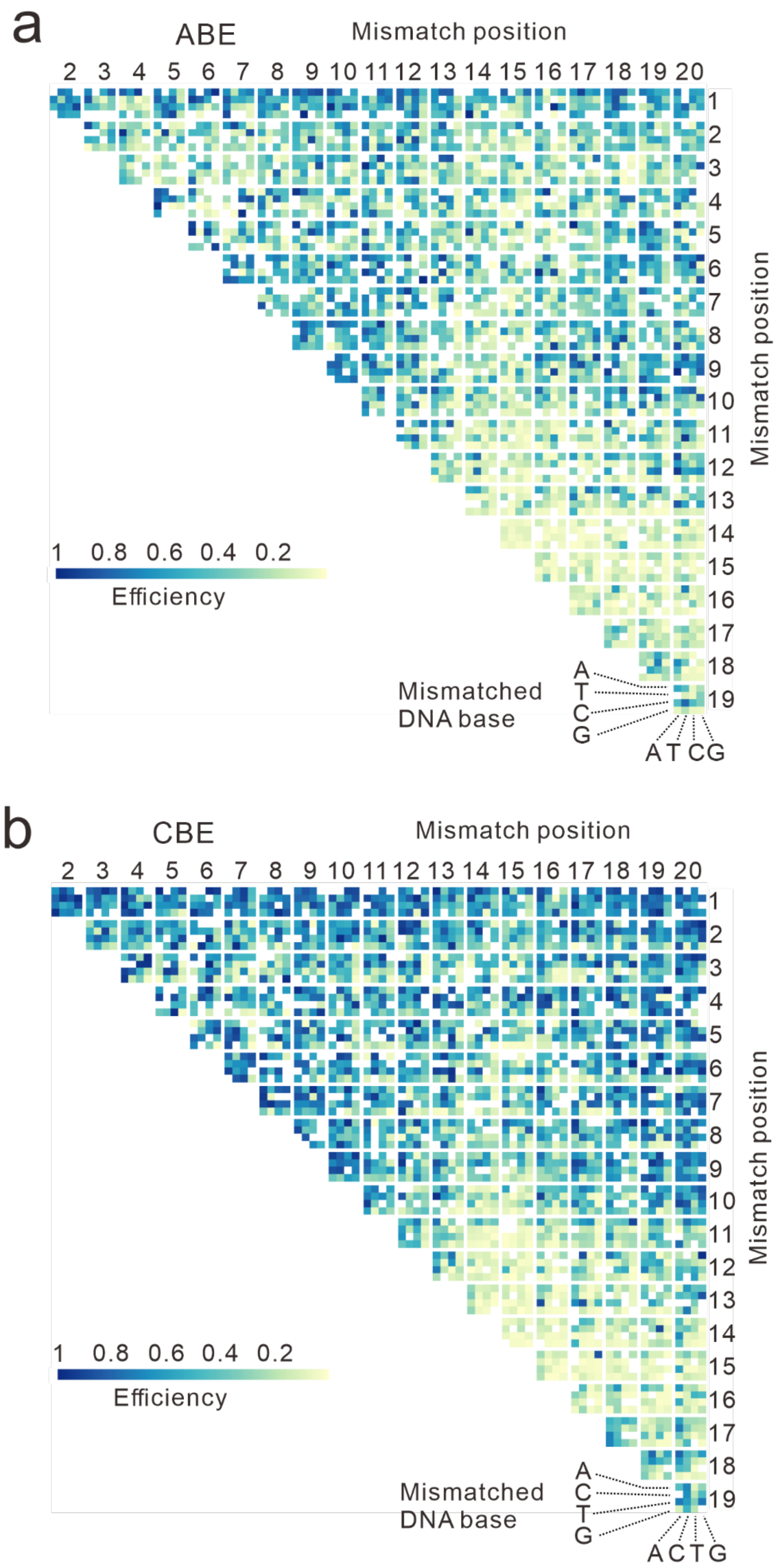


**Supplementary Figure 9. Positional effects of two nucleotide mismatches on conversion efficiency.**
(**a**) Influence of two nucleotide mismatches on conversion efficiency for ABE.
(**b**) Influence of two nucleotide mismatches on conversion efficiency for CBE.

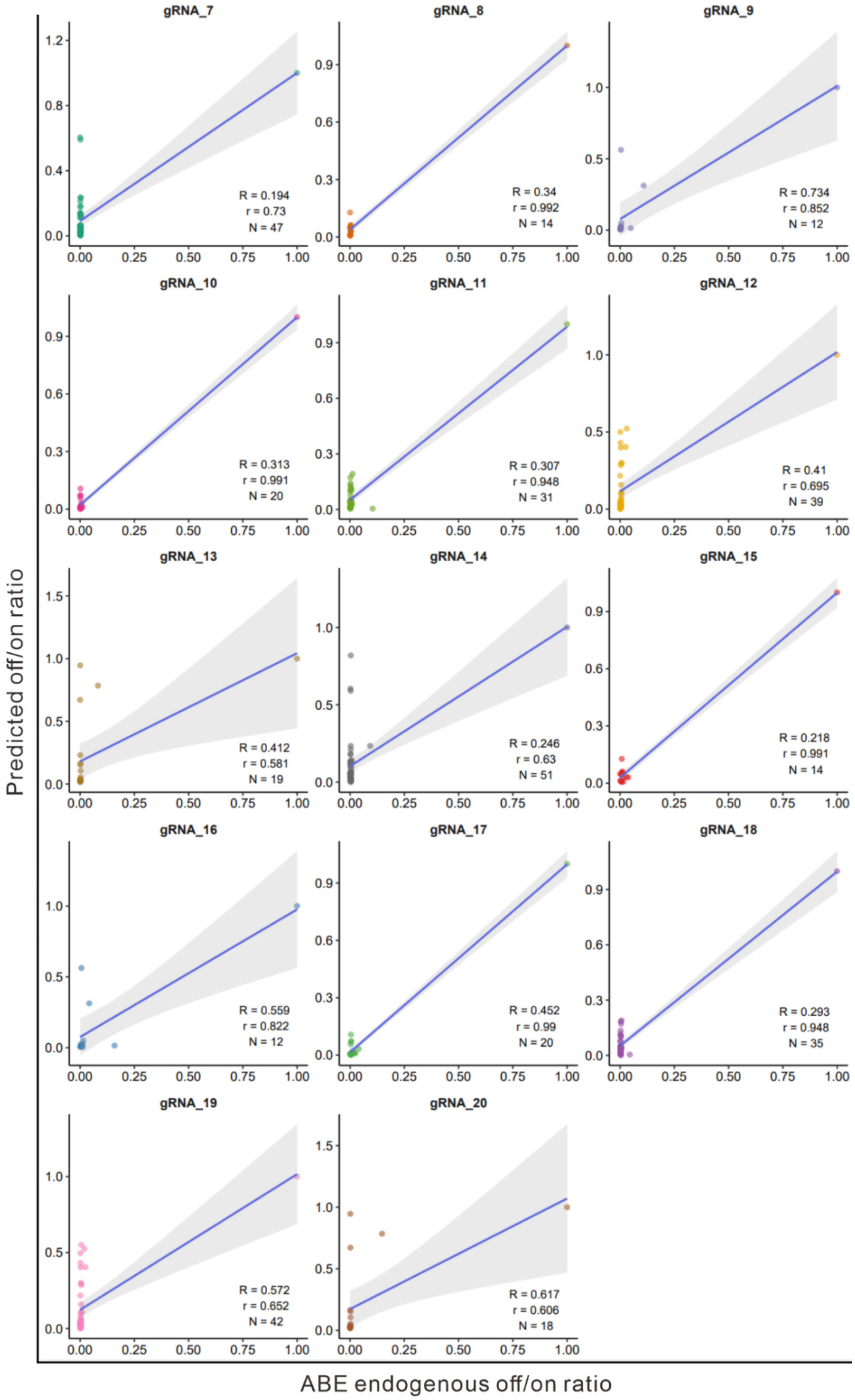


**Supplementary Figure 10. Evaluation of ABEdeepoff prediction for conversion efficiency with 14 groups of endogenous off-target datasets.**

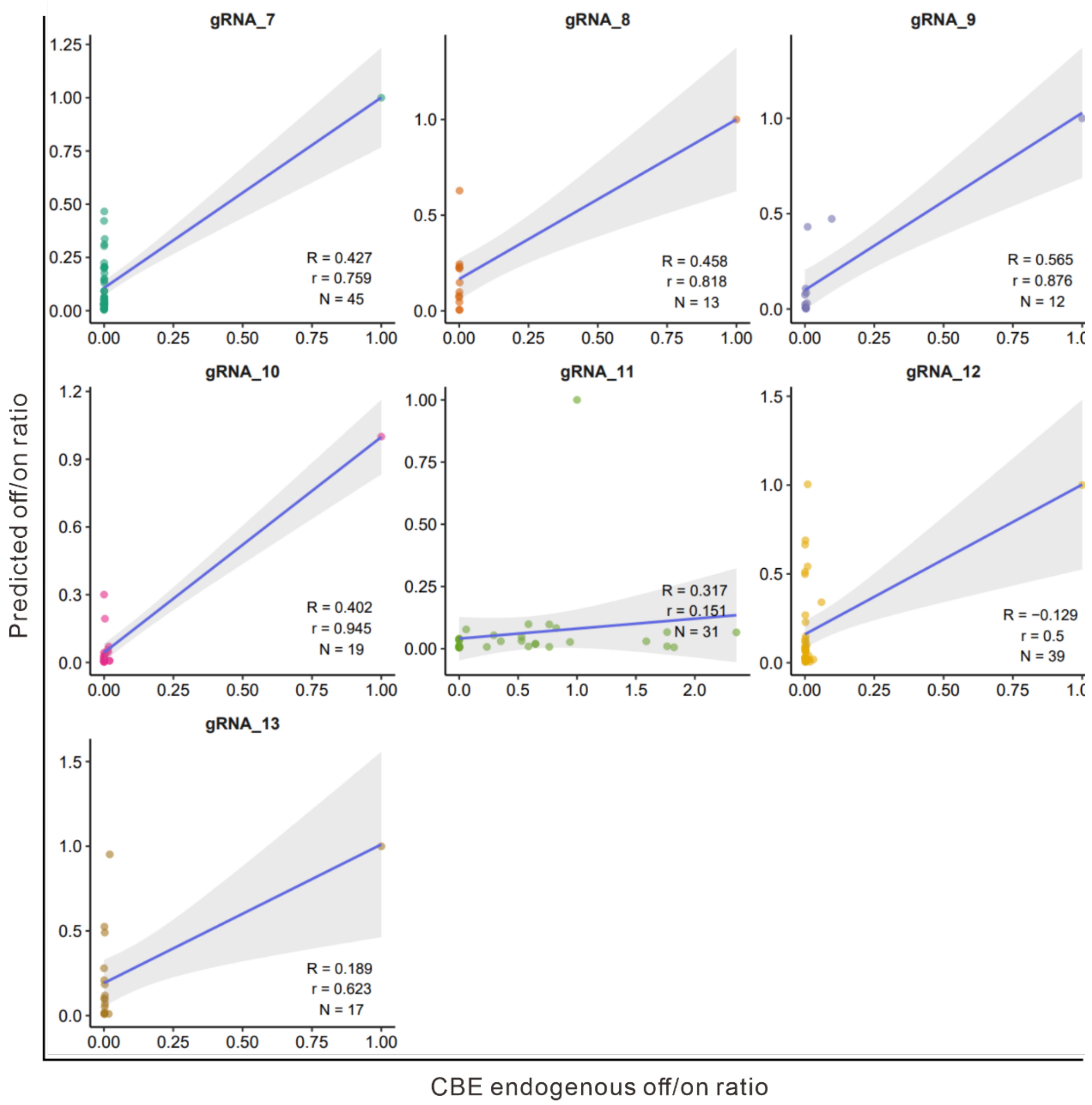


**Supplementary Figure 11. Evaluation of CBEdeepoff prediction for conversion efficiency with 7 groups of endogenous off-target datasets.**

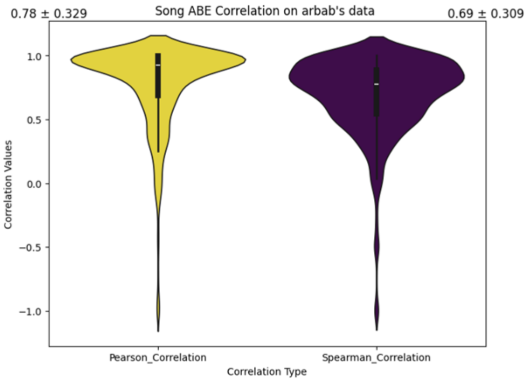
0.78 ± 0.329
Song ABE Correlation on arbab's data
0.69 ± 0.309
Correlation Values
Pearson_Correlation
Spearman_Correlation
Correlation Type

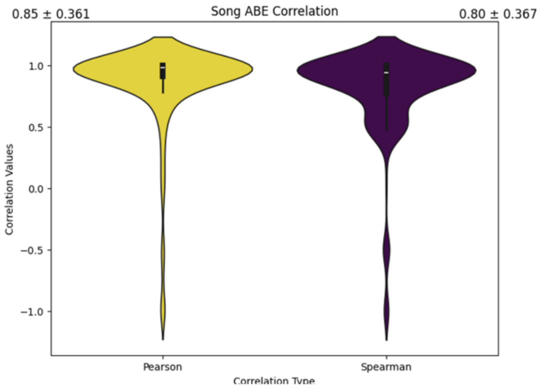
0.85 ± 0.361
Song ABE Correlation
0.80 ± 0.367
Correlation Values
Pearson
Spearman
Correlation Type

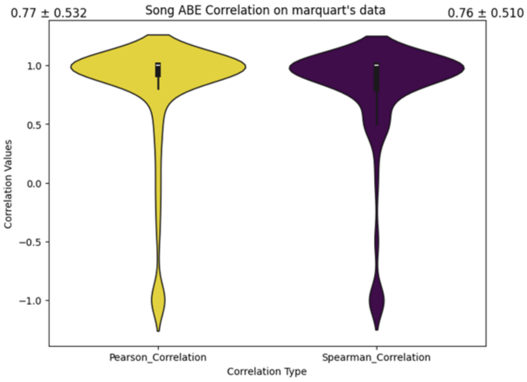
0.77 ± 0.532
Song ABE Correlation on marquart's data
0.76 ± 0.510
Correlation Values
Pearson_Correlation
Spearman_Correlation
Correlation Type

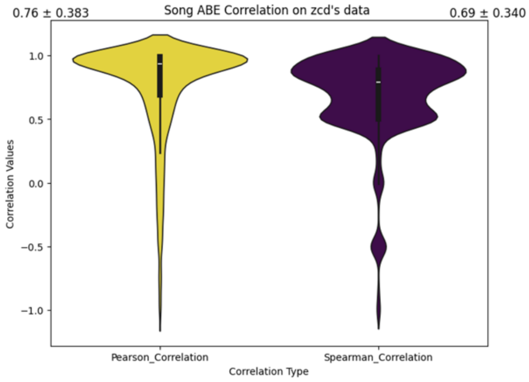
0.76 ± 0.383
Song ABE Correlation on zcd's data
0.69 ± 0.340
Correlation Values
Pearson_Correlation
Spearman_Correlation
Correlation Type

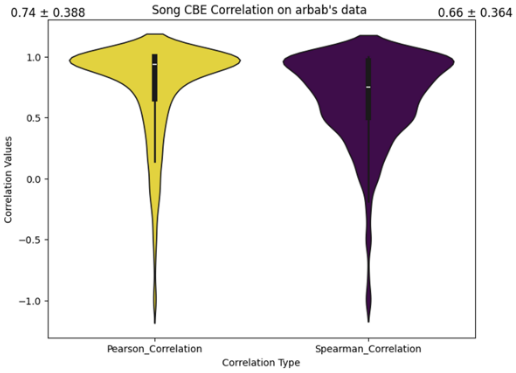
0.74 ± 0.388
Song CBE Correlation on arbab's data
0.66 ± 0.364
Correlation Values
Pearson_Correlation
Spearman_Correlation
Correlation Type

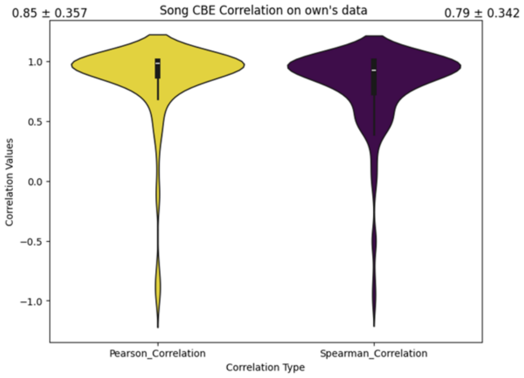
0.85 ± 0.357
Song CBE Correlation on own's data
0.79 ± 0.342
Correlation Values
Pearson_Correlation
Spearman_Correlation
Correlation Type

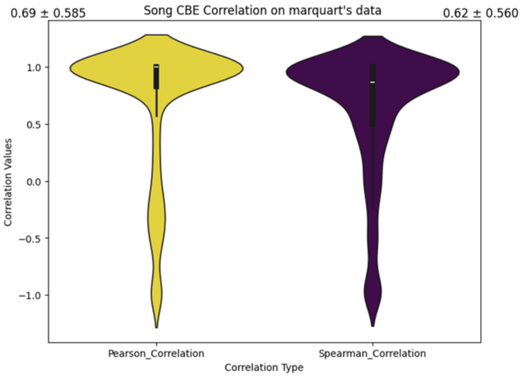
0.69 ± 0.585
Song CBE Correlation on marquart's data
0.62 ± 0.560
Correlation Values
Pearson_Correlation
Spearman_Correlation
Correlation Type

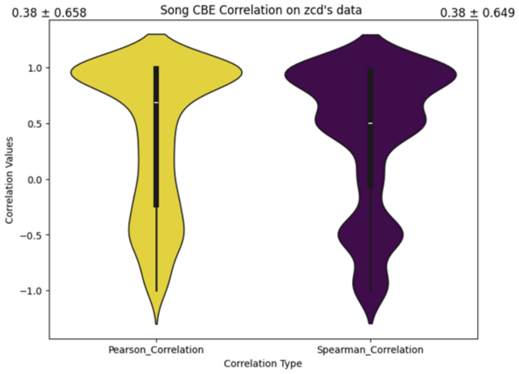
0.38 ± 0.658
Song CBE Correlation on zcd's data
0.38 ± 0.649
Correlation Values
Pearson_Correlation
Spearman_Correlation
Correlation Type

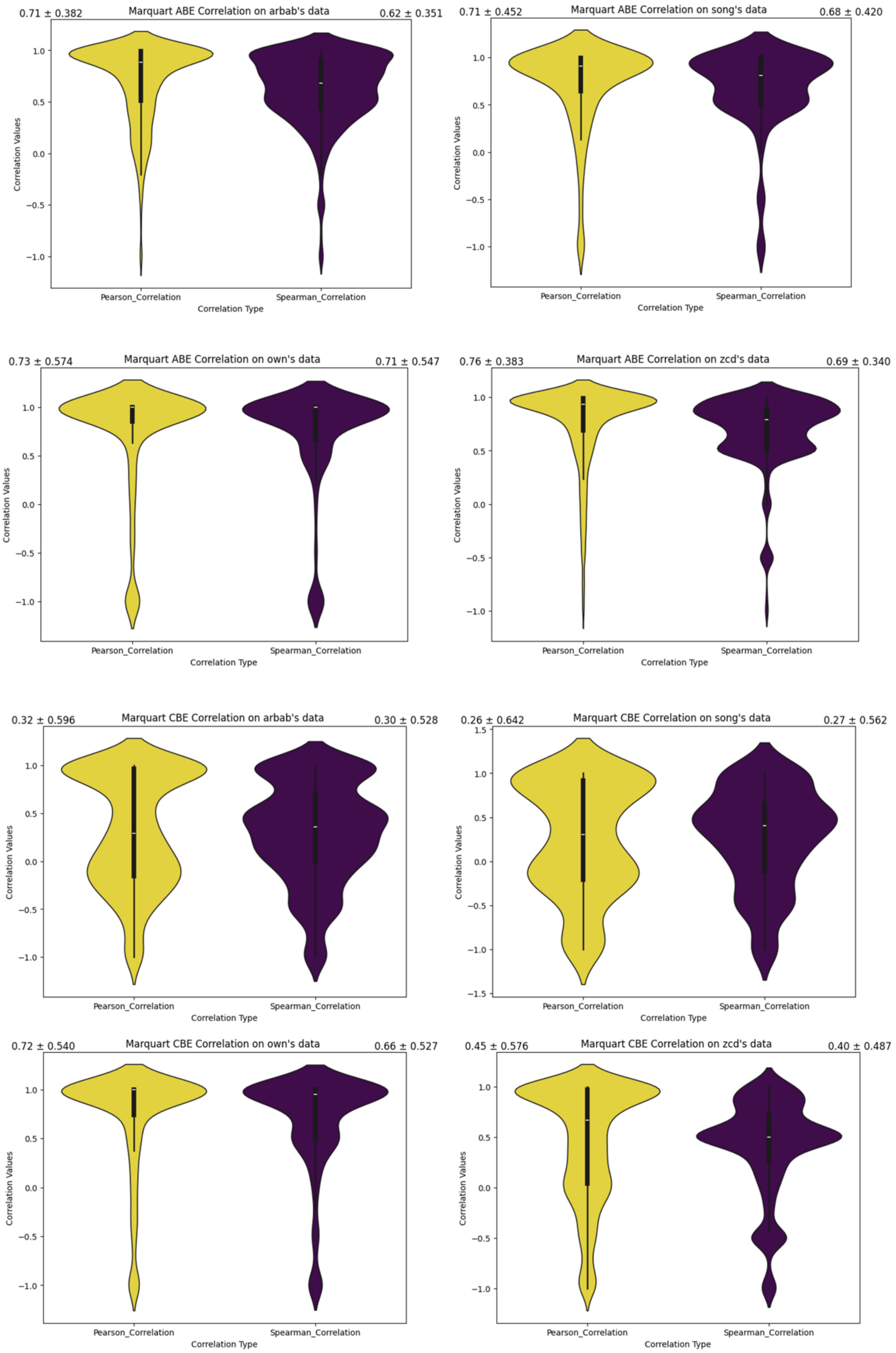
0.71 ± 0.382
Marquart ABE Correlation on arbab's data
0.62 ± 0.351
0.71 ± 0.452
Marquart ABE Correlation on song's data
0.68 ± 0.420
0.73 ± 0.574
Marquart ABE Correlation on own's data
0.71 ± 0.547
0.76 ± 0.383
Marquart ABE Correlation on zcd's data
0.69 ± 0.340
0.32 ± 0.596
Marquart CBE Correlation on arbab's data
0.30 ± 0.528
0.26 ± 0.642
Marquart CBE Correlation on song's data
0.27 ± 0.562
0.72 ± 0.540
Marquart CBE Correlation on own's data
0.66 ± 0.527
0.45 ± 0.576
Marquart CBE Correlation on zcd's data
0.40 ± 0.487
Correlation Values
Pearson_Correlation
Spearman_Correlation
Correlation Type

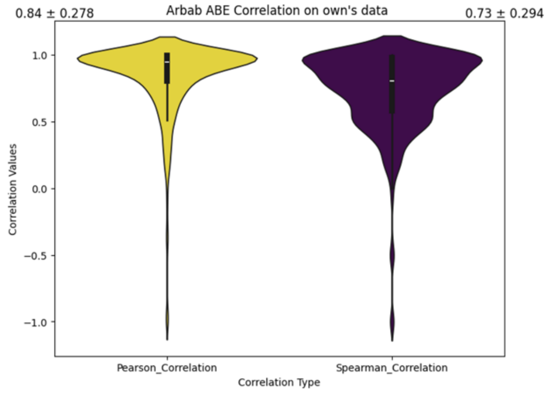
0.84 ± 0.278
Arbab ABE Correlation on own's data
0.73 ± 0.294
Correlation Values
1.0
0.5
0.0
−0.5
−1.0
Pearson_Correlation
Spearman_Correlation
Correlation Type

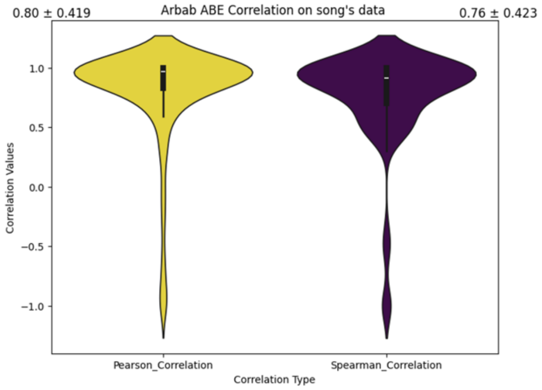
0.80 ± 0.419
Arbab ABE Correlation on song's data
0.76 ± 0.423
Correlation Values
1.0
0.5
0.0
−0.5
−1.0
Pearson_Correlation
Spearman_Correlation
Correlation Type

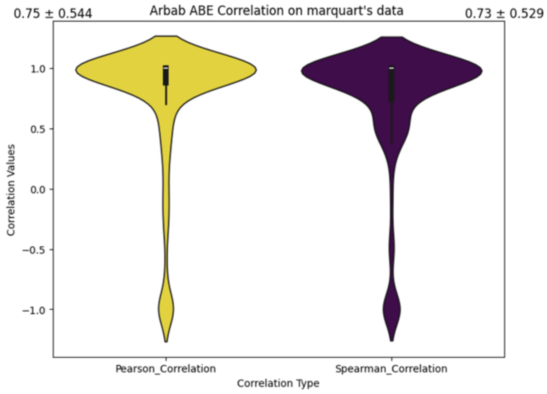
0.75 ± 0.544
Arbab ABE Correlation on marquart's data
0.73 ± 0.529
Correlation Values
1.0
0.5
0.0
−0.5
−1.0
Pearson_Correlation
Spearman_Correlation
Correlation Type

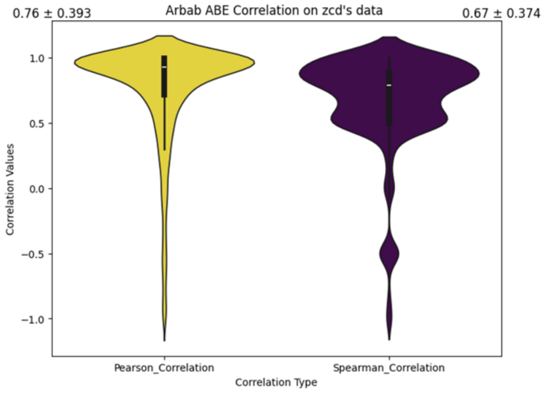
0.76 ± 0.393
Arbab ABE Correlation on zcd's data
0.67 ± 0.374
Correlation Values
1.0
0.5
0.0
−0.5
−1.0
Pearson_Correlation
Spearman_Correlation
Correlation Type

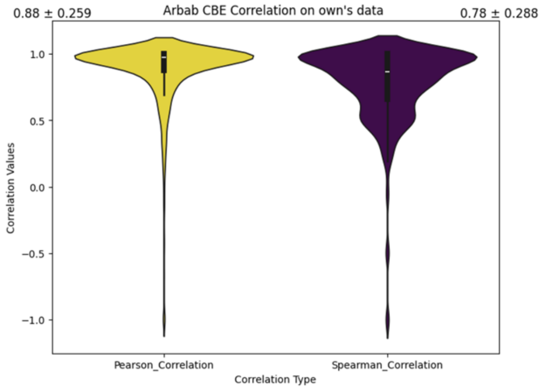
0.88 ± 0.259
Arbab CBE Correlation on own's data
0.78 ± 0.288
Correlation Values
1.0
0.5
0.0
−0.5
−1.0
Pearson_Correlation
Spearman_Correlation
Correlation Type

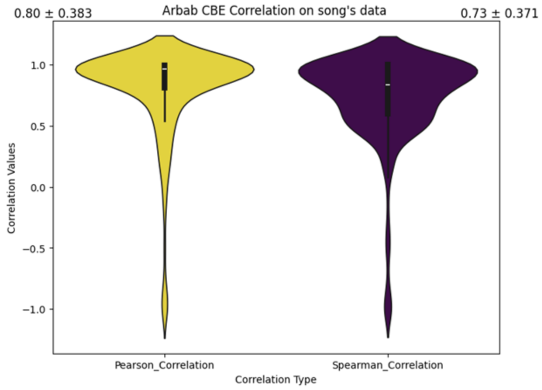
0.80 ± 0.383
Arbab CBE Correlation on song's data
0.73 ± 0.371
Correlation Values
1.0
0.5
0.0
−0.5
−1.0
Pearson_Correlation
Spearman_Correlation
Correlation Type

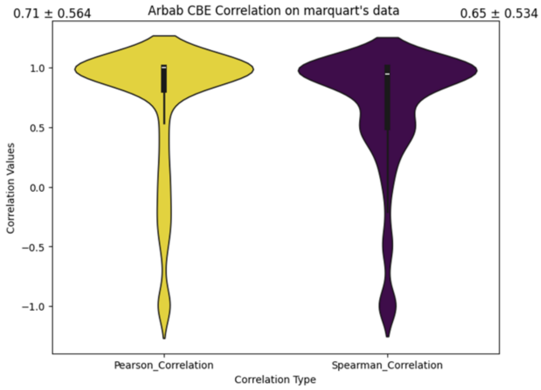
0.71 ± 0.564
Arbab CBE Correlation on marquart's data
0.65 ± 0.534
Correlation Values
1.0
0.5
0.0
−0.5
−1.0
Pearson_Correlation
Spearman_Correlation
Correlation Type

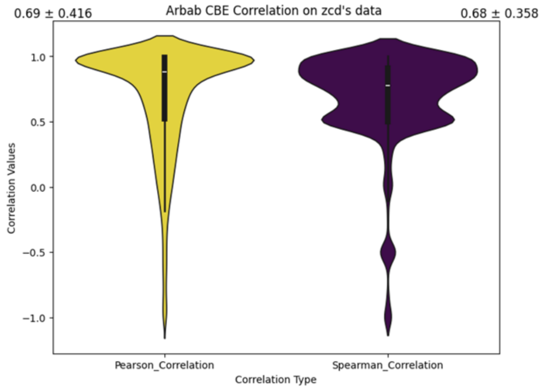
0.69 ± 0.416
Arbab CBE Correlation on zcd's data
0.68 ± 0.358
Correlation Values
1.0
0.5
0.0
−0.5
−1.0
Pearson_Correlation
Spearman_Correlation
Correlation Type

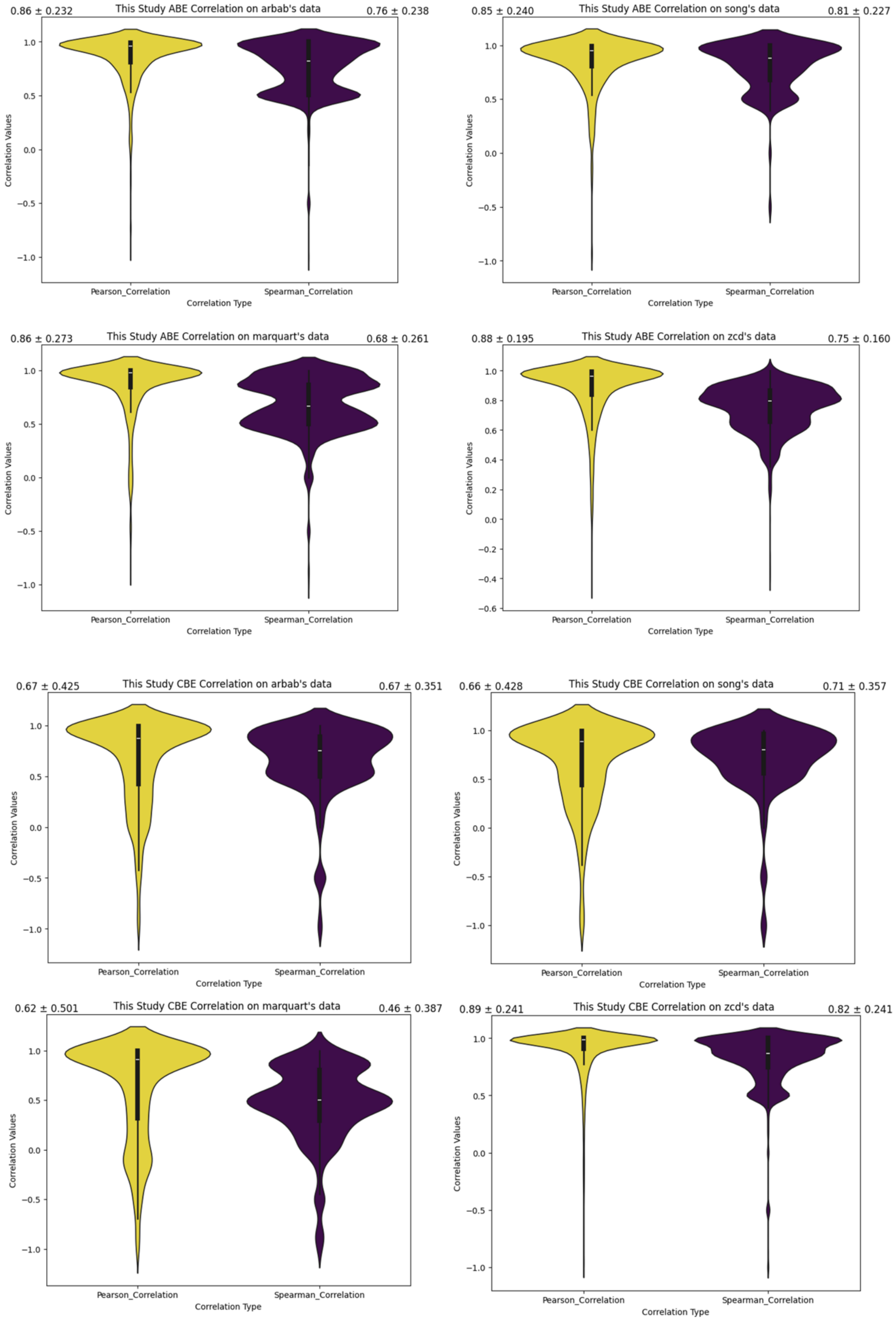


**Supplementary Figure 12. Benchmarking of Bedeepon with other four models via mean Spearman and Pearson Correlation with variance on edited outcomes.**

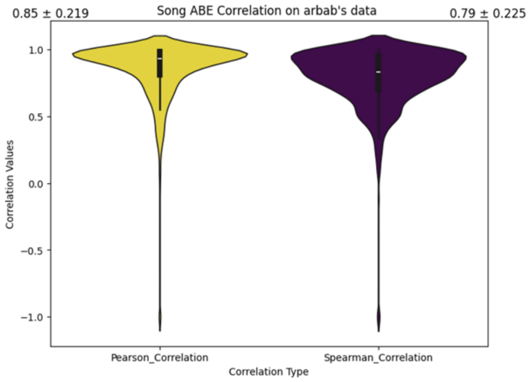
0.85 ± 0.219
Song ABE Correlation on arbab's data
0.79 ± 0.225
Correlation Values
Pearson_Correlation
Spearman_Correlation
Correlation Type

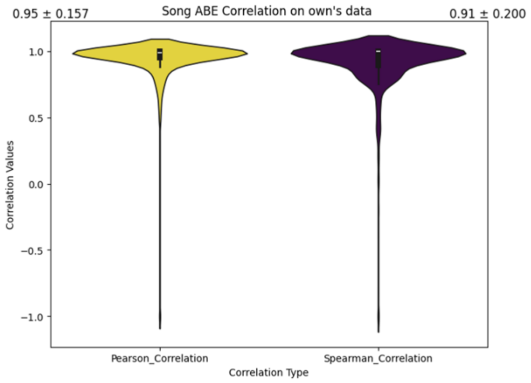
0.95 ± 0.157
Song ABE Correlation on own's data
0.91 ± 0.200
Correlation Values
Pearson_Correlation
Spearman_Correlation
Correlation Type

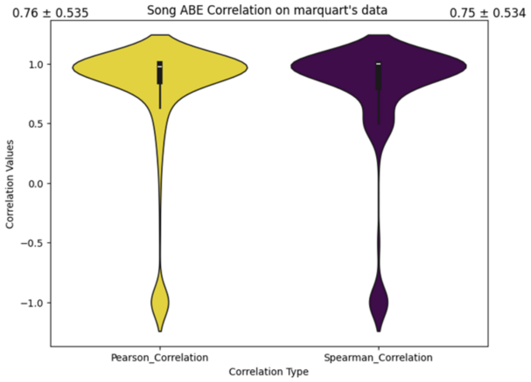
0.76 ± 0.535
Song ABE Correlation on marquart's data
0.75 ± 0.534
Correlation Values
Pearson_Correlation
Spearman_Correlation
Correlation Type

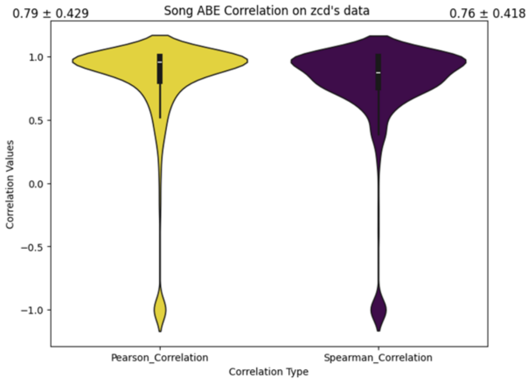
0.79 ± 0.429
Song ABE Correlation on zcd's data
0.76 ± 0.418
Correlation Values
Pearson_Correlation
Spearman_Correlation
Correlation Type

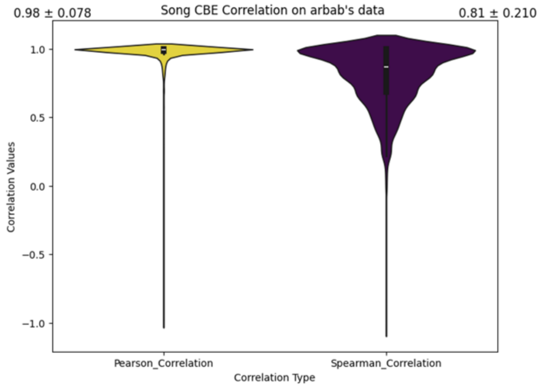
0.98 ± 0.078
Song CBE Correlation on arbab's data
0.81 ± 0.210
Correlation Values
Pearson_Correlation
Spearman_Correlation
Correlation Type

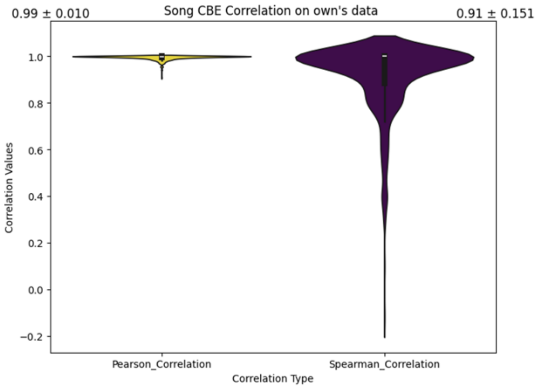
0.99 ± 0.010
Song CBE Correlation on own's data
0.91 ± 0.151
Correlation Values
Pearson_Correlation
Spearman_Correlation
Correlation Type

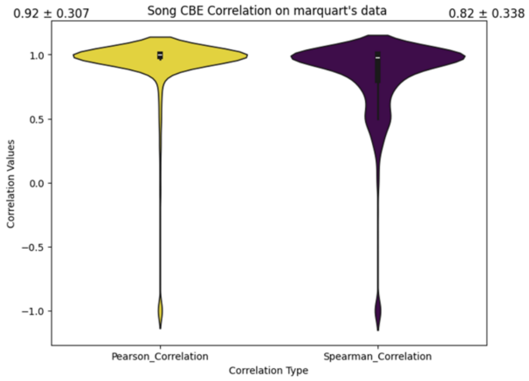
0.92 ± 0.307
Song CBE Correlation on marquart's data
0.82 ± 0.338
Correlation Values
Pearson_Correlation
Spearman_Correlation
Correlation Type

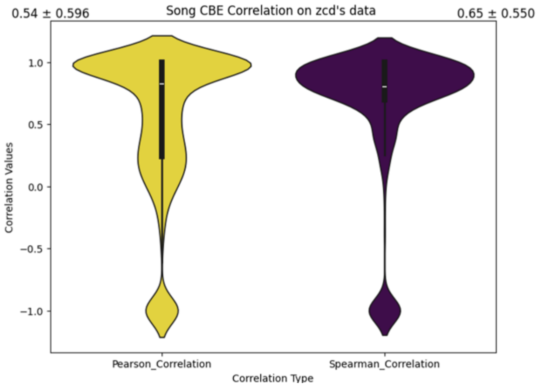
0.54 ± 0.596
Song CBE Correlation on zcd's data
0.65 ± 0.550
Correlation Values
Pearson_Correlation
Spearman_Correlation
Correlation Type

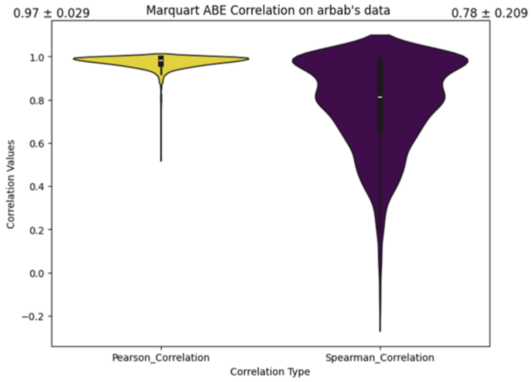
0.97 ± 0.029
Marquart ABE Correlation on arbab's data
0.78 ± 0.209
Correlation Values
Pearson_Correlation
Spearman_Correlation
Correlation Type

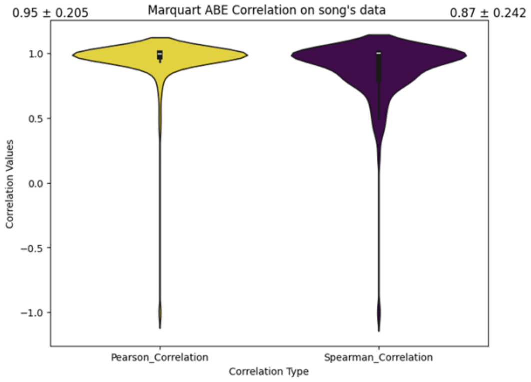
0.95 ± 0.205
Marquart ABE Correlation on song's data
0.87 ± 0.242
Correlation Values
Pearson_Correlation
Spearman_Correlation
Correlation Type

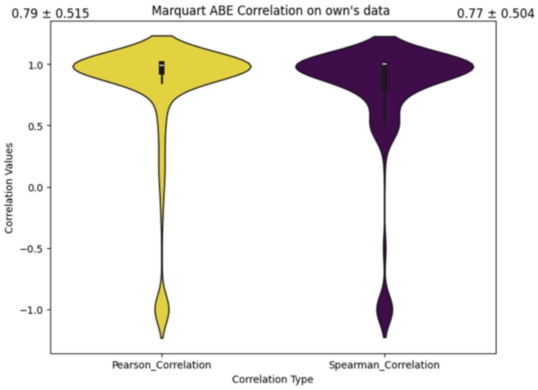
0.79 ± 0.515
Marquart ABE Correlation on own's data
0.77 ± 0.504
Correlation Values
Pearson_Correlation
Spearman_Correlation
Correlation Type

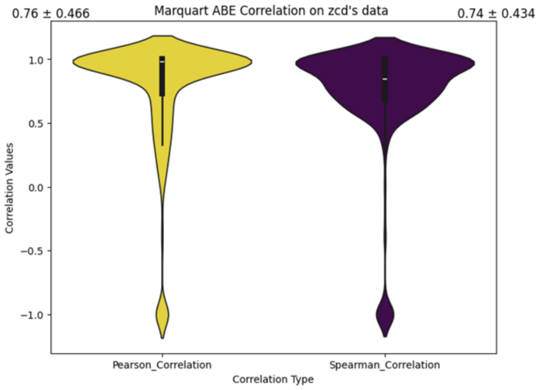
0.76 ± 0.466
Marquart ABE Correlation on zcd's data
0.74 ± 0.434
Correlation Values
Pearson_Correlation
Spearman_Correlation
Correlation Type

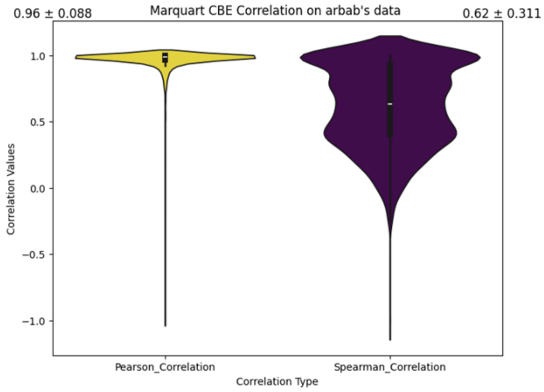
0.96 ± 0.088
Marquart CBE Correlation on arbab's data
0.62 ± 0.311
Correlation Values
Pearson_Correlation
Spearman_Correlation
Correlation Type

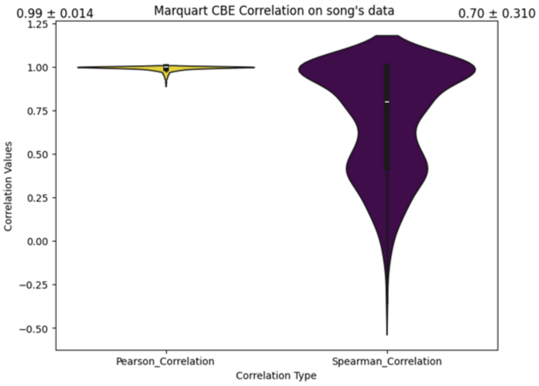
0.99 ± 0.014
Marquart CBE Correlation on song's data
0.70 ± 0.310
Correlation Values
Pearson_Correlation
Spearman_Correlation
Correlation Type

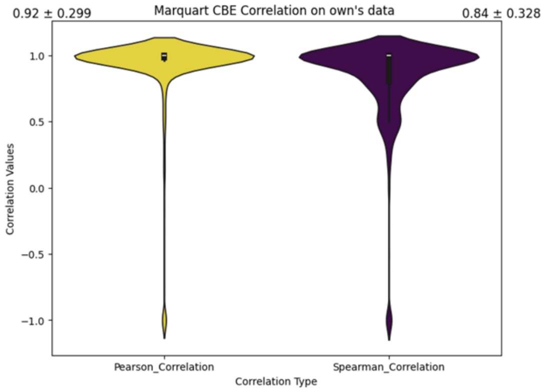
0.92 ± 0.299
Marquart CBE Correlation on own's data
0.84 ± 0.328
Correlation Values
Pearson_Correlation
Spearman_Correlation
Correlation Type

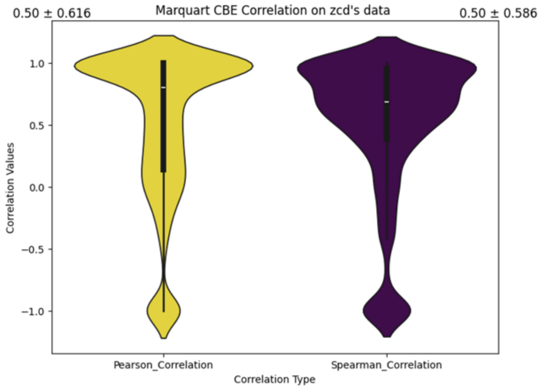
0.50 ± 0.616
Marquart CBE Correlation on zcd's data
0.50 ± 0.586
Correlation Values
Pearson_Correlation
Spearman_Correlation
Correlation Type

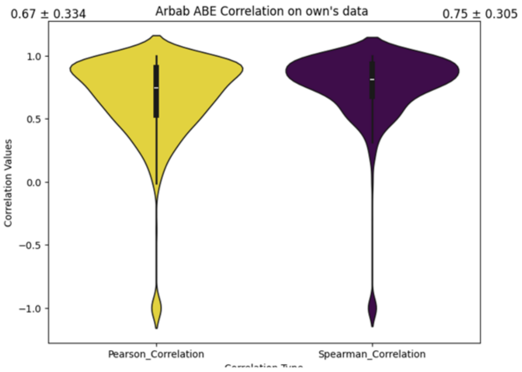
0.67 ± 0.334
Arbab ABE Correlation on own's data
0.75 ± 0.305
Correlation Values
1.0
0.5
0.0
−0.5
−1.0
Pearson_Correlation
Spearman_Correlation

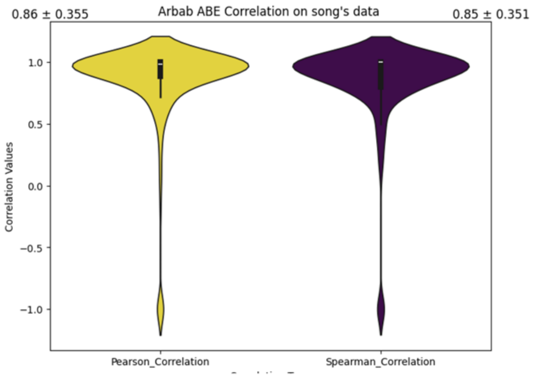
0.86 ± 0.355
Arbab ABE Correlation on song's data
0.85 ± 0.351
Correlation Values
1.0
0.5
0.0
−0.5
−1.0
Pearson_Correlation
Spearman_Correlation

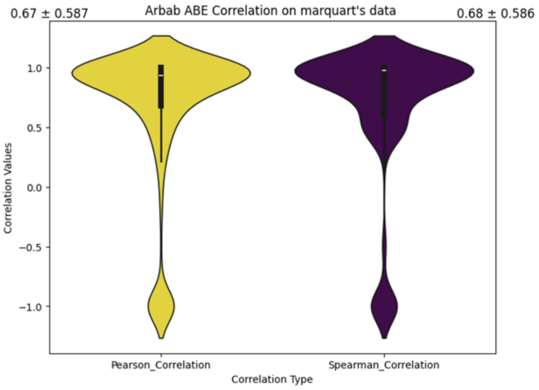
0.67 ± 0.587
Arbab ABE Correlation on marquart's data
0.68 ± 0.586
Correlation Values
1.0
0.5
0.0
−0.5
−1.0
Pearson_Correlation
Spearman_Correlation
Correlation Type

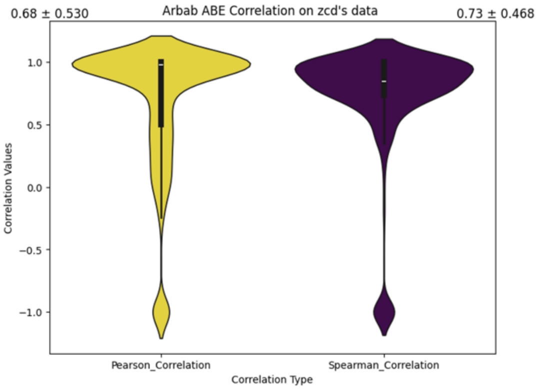
0.68 ± 0.530
Arbab ABE Correlation on zcd's data
0.73 ± 0.468
Correlation Values
1.0
0.5
0.0
−0.5
−1.0
Pearson_Correlation
Spearman_Correlation
Correlation Type

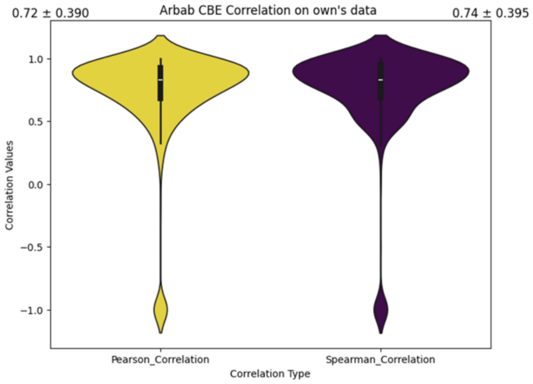
0.72 ± 0.390
Arbab CBE Correlation on own's data
0.74 ± 0.395
Correlation Values
1.0
0.5
0.0
−0.5
−1.0
Pearson_Correlation
Spearman_Correlation
Correlation Type

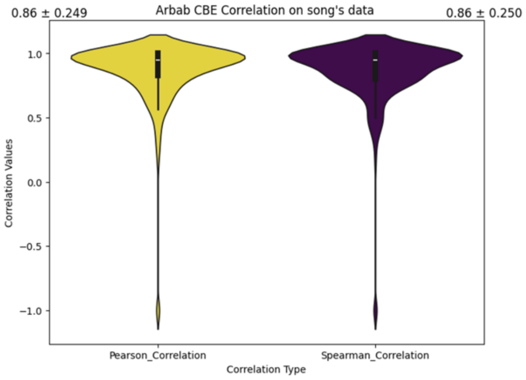
0.86 ± 0.249
Arbab CBE Correlation on song's data
0.86 ± 0.250
Correlation Values
1.0
0.5
0.0
−0.5
−1.0
Pearson_Correlation
Spearman_Correlation
Correlation Type

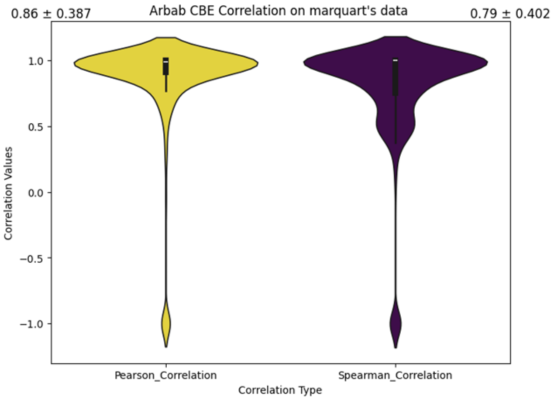
0.86 ± 0.387
Arbab CBE Correlation on marquart's data
0.79 ± 0.402
Correlation Values
1.0
0.5
0.0
−0.5
−1.0
Pearson_Correlation
Spearman_Correlation
Correlation Type

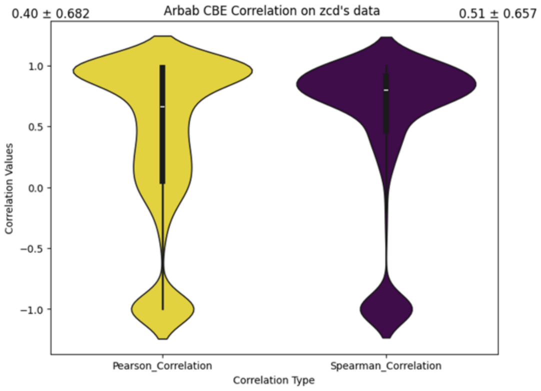
0.40 ± 0.682
Arbab CBE Correlation on zcd's data
0.51 ± 0.657
Correlation Values
1.0
0.5
0.0
−0.5
−1.0
Pearson_Correlation
Spearman_Correlation
Correlation Type

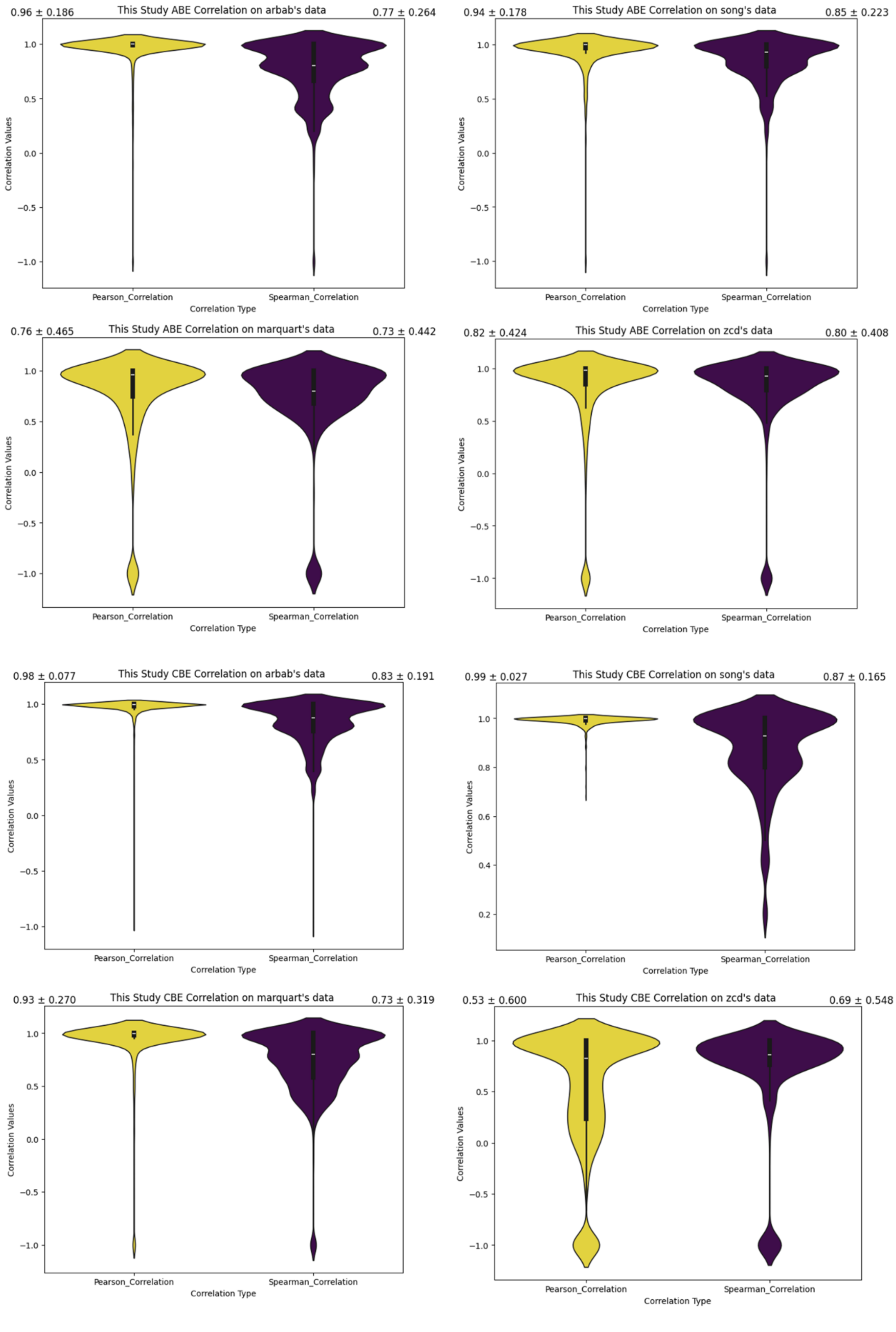


**Supplementary Figure 13. Benchmarking of Bedeepon with other four models via mean Spearman and Pearson Correlation with variance on overall outcomes.**

**Supplementary Table：**

Supplementary Table 1.The efficiencies of base editors in 293T and Hela cell lines (provided as a separate excel file).
Supplementary Table 2. The outcomes dataset of 60,615 gRNAs for ABE in HEK293T cells (DOIs: https://doi.org/10.6084/m9.figshare.13525568.v2 ).
Supplementary Table 3. The outcomes dataset of 73,303 gRNAs for CBE in HEK293T cells (DOIs: https://doi.org/10.6084/m9.figshare.13525568.v2 ).
Supplementary Table 4. The outcomes dataset of 58,445 gRNAs for ABE in HeLa cells (DOIs: https://doi.org/10.6084/m9.figshare.13525568.v2 ).
Supplementary table 5. The editing efficiency and outcomes of endogenous loci in iPSCs (provided as a separate excel file).
Supplementary Table 6. The outcomes dataset of 56,529 gRNAs for CBE in HeLa cells (DOIs: https://doi.org/10.6084/m9.figshare.13525568.v2 ).
Supplementary Table 7. The designed off-target oligonucleotide library for ABE and CBE (provided as a separate excel file).
Supplementary Table 8. The efficiency dataset of mutation targets for ABE (provided as a separate excel file).
Supplementary Table 9. The efficiency dataset of mutation targets for CBE.xlsx
Supplementary Table 10. The third-party off target datasets of ABE (provided as a separate excel file).
Supplementary Table 11. The third-party off target datasets of CBE (provided as a separate excel file).
Supplementary Table 12. The sequences of oligonucleotides (provided as a separate excel file).
Supplementary Table 13. Benchmarking scores of four models (provided as a separate excel file).
